# Peptide-PAINT using a transfected-docker enables live- and fixed-cell super-resolution imaging

**DOI:** 10.1101/2022.09.07.507019

**Authors:** Barun Kumar Maity, Duncan Nall, Yongjae Lee, Paul R Selvin

## Abstract

Point accumulation for imaging in nanoscale topography (PAINT) is a single-molecule technique for super-resolution microscopy, achieving ∼5-25 nanometer resolution. Here we show that by transfecting the protein-of-interest with a docker-coil, rather than by adding the docker externally—as is the norm when using DNA tethers or antibodies as dockers—we can achieve similar localization, ∼10 nm. However, using a transfected docker has several experimental advances and simplifications. Most importantly, it allows Peptide-PAINT to be applied to transfected *live* cells, including surface proteins in mammalian cells and neurons under physiological conditions. The enhance resolution of Peptide-PAINT is also shown for organelles in fixed cells to unravel structural details including ≈40-nm and ≈60-nm axial repeats in vimentin filaments in the cytoplasm, and fiber shapes of sub-100-nm histone-rich regions in the nucleus.

## INTRODUCTION

Single-molecule super-resolution microscopy can visualize structural details in life sciences beyond the ∼250 nm diffraction limited resolution, achieving 5-25 nm resolution^1–4^. These techniques have been implemented in several ways, including stochastic optical reconstruction microscopy (STORM)^1^, fluorescence photoactivated localization microscopy ((F)PALM)^2,5^, point accumulation for imaging in nanoscale topography (PAINT)^6^, as well as stimulated emission depletion (STED) microscopy ^7^. Except for STED, these super-resolution techniques require that only one fluorophore emits at a time. With a single molecule, the average position of the fluorophore can be localized to ∼ λ/2N^1/2^ (λ: emission wavelength, N: total number of emitted photons). STORM and PALM do this by utilizing stochastic blinking of individual fluorophores, often requiring additional chemicals or lasers. This can limit the type of systems that can be examined. In contrast, PAINT does not require these^3,4,8–10^. Rather, PAINT achieves single molecule blinking by transient and repetitive binding of a fluorescently labeled imager strand to a docker strand that is appended to the target-of-interest. The imager produces a transient high efflux of localized fluorescent photons while hybridized, enabling localization of a single target site with nanometer accuracy while discriminating against the background created by the fast diffusing non-hybridized imager strands^4^.

PAINT provides some advantages over STORM and PALM. These include: no need for a blinking dye or photoactivable fluorescent proteins; no additional chemicals in the imaging buffer to make the fluorophores blink; less photo-bleaching due to continuous exchange of imager; better spatial resolution (at a slower rate of data acquisition); and high multiplexing ability^4,8,9,11^. PAINT can also determine the relative or absolute stoichiometry of the underlying docker concentration based on binding-unbinding kinetics. This kinetic scheme is called quantitative PAINT (qPAINT) and is generally not straightforward with STORM/PALM because of the stochastic nature of the blinking phenomenon.^12–14^

While DNA oligos are commonly used in PAINT, peptides that hybridize to each other have recently been used (named Peptide-PAINT).^10,15–19^ For example, Fischer et al. used a fluorescently labeled imager peptide in fixed cells to transiently bind to the extracellular side of talin, a protein involved in cell adhesion process.^16^ This particular system is highly advantageous as the protein-of-interest (talin) already has a binding site for the imager, and hence there is no need to have an external docker-peptide.

Another example of Peptide-PAINT is SYNZIP17-SYNZIP18, a coiled-coil peptide pair that was employed for imaging *intracellular* targets in live yeast cells where both the docker, genetically appended to the target-of-interest, and the imager, conjugated to a fluorescent protein (FP ∼25kD), were expressed inside cells^15^. This is the only case of Peptide-PAINT on living cells (as far as we know). This, however, was not shown to work on mammalian cells and the use of FPs is limiting because of their relatively weak fluorescence, poor photostability and large size.

Like the SYNZIP17-SYNZIP18 pair, a FP (mNeonGreen) was used by Tas et al. as the imager to transiently bind to a small docker (DTWV) appended to Erbin PDS (MW∼10k). It was optimized on fixed cells, and hence is unknown if it will work in a live cell. It also has the disadvantages of using a relatively large protein (an FP) as the imager. In contrast, using a bright (organic) fluorophore, and a modified form of the heterodimeric E/K peptide pair-E22 and CK19, Eklund et al. demonstrated super-resolution imaging of filament proteins on fixed cells^10^. In addition, the binding rate of this pair (k_on_) in an *in vitro* experiment, i.e., on a coverslip, is ∼388 fold faster than the result of Tas et al. (who used a FP containing imager).^33^ However, Eklund et al. attached the E22-docker externally to the protein-of-interest via an antibody, instead of transfecting the docker coil with the protein-of-interest.

In this report, we report nanometer resolution with PAINT between a genetically expressed E22-docker to an externally expressed CK19 imager of various targets in fixed and live HeLa cells and importantly, to a live neuron in physiological buffer conditions. This establishes the first case of Peptide-PAINT imaging on live mammalian cells. The work highlights practical advantage of Peptide-PAINT imaging in fixed cells and applicability to surface proteins in live cells.

## RESULTS AND DISCUSSIONS

The overall labeling scheme is shown in Fig. 1A, where we show Peptide-PAINT for the two proteins, vimentin (Fig. 1B-H) and Golgi body (Fig. 1I-L). In both cases, we attached a GFP and a C-terminal E22 docker (i.e., protein-GFP-E22). After transfection, and subsequent fixation and permeabilization in a HeLa cell, the GFP fluorescence identifies the transfected cells and enables a diffraction-limited image of the target proteins (Fig. 1B for vimentin and 1I for Golgi body). The CK19 imager was coupled to the dye LD655 and was used to obtain a high-resolution image by transiently hybridizing to the E22 docker via Peptide-PAINT (for details, see Methods). As expected, we observed fluorescence blinking from transient hybridization of the imager. This does not occur when the cells were transfected with the plasmid *without* the docker sequence (for Vimentin-GFP, see Fig. S1C and S1F). These results indicate that the observed blinking is due to specific transient binding-unbinding between the docker and the imager. Several such binding events were localized with high precision and a super-resolution image was constructed (Fig. 1C for vimentin, 1J for Golgi body). For vimentin (Golgi bodies), the mean photon counts are 614 (723) photons (Fig. S2A and S2B), and the localization precisions are 15 (14) nm (Fig. S2D and S2E), analyzed using Picasso software package^21^ (Localization precisions are approximately a factor of 1.5-2 larger when analyzed using ThunderSTORM, another commonly used software package^22^: see SI, Table-1, Methods). The magnified reconstructed images of Golgi body (Fig. 1L) clearly show the well-resolved structures compared to their diffraction-limited ones (Fig. 1K). This establishes that Peptide-PAINT imaging with directly appended docker can improve the resolution significantly greater than the diffraction limit of ∼250 nm.

**Figure 1:**
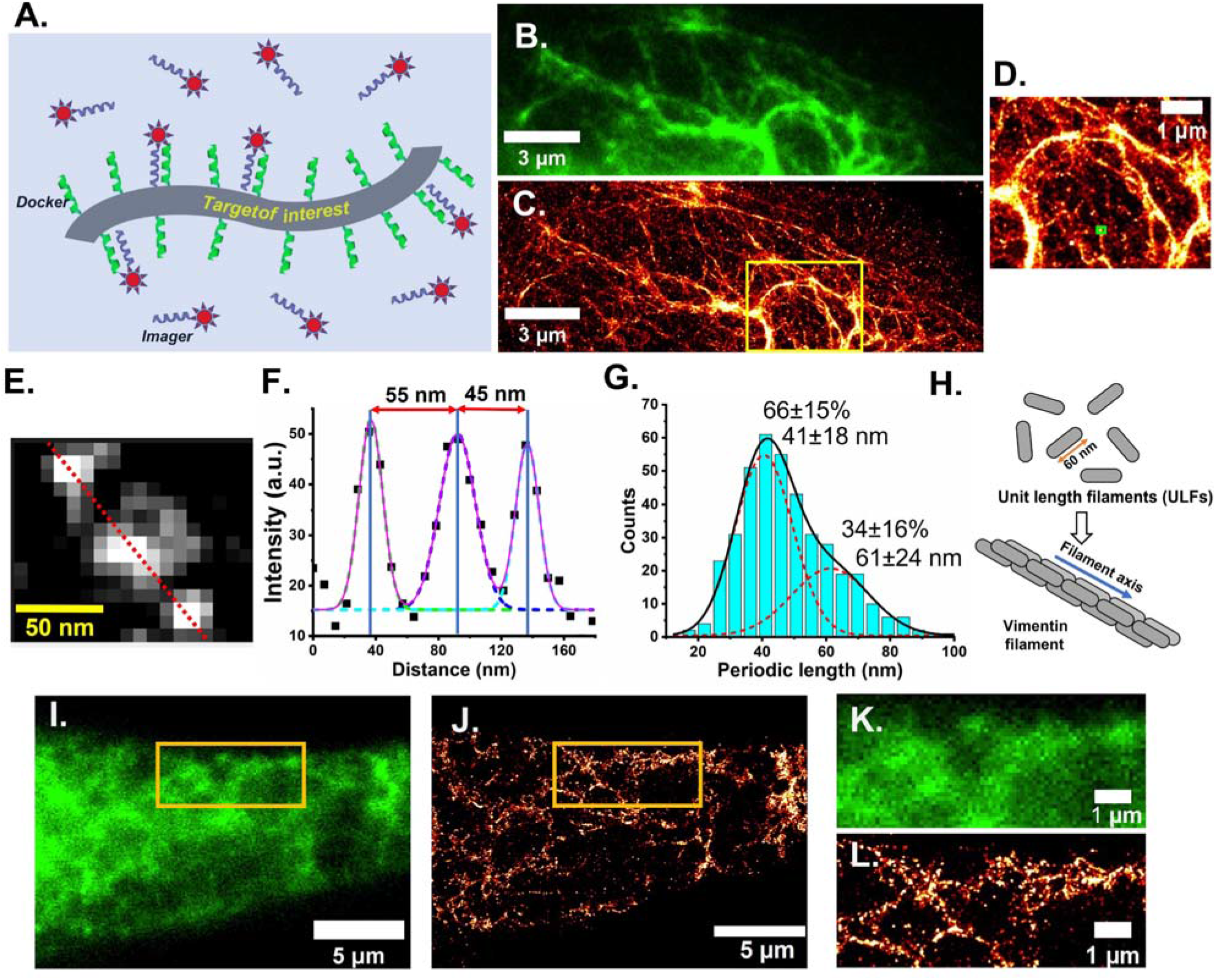
Peptide-PAINT and Diffraction-limited imaging of vimentin and Golgi bodies. (A) Schematic of the experiment. All samples were on fixed HeLa cells with the vimentin (B-H) or Golgi bodies (I-L) labeled with GFP and E22 docker. In each case, diffraction limited images were obtained by using GFP fluorescence, and super-resolution images by Peptide-PAINT using 0.5 nM LD655-CK19 in PBS supplemented with 2% BSA as imager. (B, C) Representative diffraction-limited and super-resolution images of vimentin. (D) Magnified image of yellow rectangular region in 1C. (E) Representative magnified image of green rectangular box in Fig. 1D representing single vimentin filament. (F) Intensity profile along the red-dotted line in Fig. 1E. (G) Periodic length distribution of vimentin filament; (H) Cartoon of vimentin filament formation. (I, J) Represent diffraction-limited and super-resolution images of Golgi body, respectively. (K, L): Magnified image of yellow rectangular region in Figure 1I and 1J, respectively.

We next focus on unraveling nanoscale structural details of vimentin filaments using its super-resolution image by Peptide-PAINT. It is already known that vimentin filament is formed by longitudinal self-assembly of short filament precursors, known as unit-length-filament (ULF)^23^ of length ∼60 nm, measured by *in vitro* EM study^23^. Recently, Vincente et al. have also reported similar axial repeats in vimentin in fixed cells where they found a pairwise distance between C-to-C terminal was ∼50 nm and between N-to-N terminal was ∼34 nm, measured by 2D STORM imaging. *In vitro*, they used a fixed vimentin-Y117L mutant, which stops polymerizing at ULF stage^24^. However, this study did not clearly establish whether the unmodified ULFs longitudinally assemble to form the actual vimentin filaments in cells. To examine this, we chose individual vimentin filaments (Fig. 1E) and drew the intensity profile along the filament axis (the red dotted line in Fig. 1E). The intensity distribution follows a periodic pattern. The peak-to-peak distances of the representative filament segment are 55 nm, and 45 nm respectively (Fig. 1F). Accumulated statistics, taking into account the 3-dimensional nature (the microscopy was done in 2D), suggests that there are two groups of peak-to-peak distances, one contributing 66 ± 15% at ∼41 nm (SD: 18 nm) and the other contributing 34 ± 16% of ∼61 nm (SD: 24 nm). (The number of single-filament segments in Fig. 1G is 190; of these, the number of peak-to-peak distances is 395: more representative single filament images have been shown in Fig. S3). In particular, the higher peak-to-peak distance, 61 nm, is most likely due to the ULFs in the imaging plane while the shorter distance, 41 nm, is likely due to 2D projection of three dimensionally distributed ULFs, as shown by Vincente et al.^24^. Hence, our results suggest that ULFs are axially assembled with ∼60 nm periodicity to form large vimentin filaments inside cells (1H). Furthermore, when we attach the E22 docker at the N-terminal of vimentin—i.e., HeLa cells were transfected with E22-Vimentin-GFP plasmid instead of Vimentin-GFP-E22—the peak-to-peak becomes 27 nm (SD: 7 nm) (Fig. S4C, 4D, 4E). This indicates that ULF is formed by anti-parallel assembly of vimentin monomers (Fig. S4F). Our results are consistent, but extends, the results of Vincente et al.^24^. Vincente et al. have measured the periodicity between C-to-C terminal and N-to-N terminal from ULF inside cells, but, here, we observe this periodicity in the actual filaments which is formed by the WT vimentin, not the mutant.

For histone H2B, a nuclear protein, we found that there are artifacts associated with fixation when attempting Peptide-PAINT—namely, the imager did not bind to the docker after normal fixation. To correct for this, we treated the cells with 1% sodium dodecyl sulfate (SDS) (see Method section for details). The fluorescence blinking of the LD655-CK19 imager was specific, as expected—i.e., it occurred for the cells transfected with H2B-GFP-E22, but not with H2B-GFP plasmid (Fig. S1B and S1E). We also found that taking the measurement at super-resolution yields new insights on the distribution of H2B. At the diffraction-limit, the image of GFP fluorescence shows that the H2B is generally non-homogeneously distributed in the nucleus (Fig. 2A) but that at super-resolution (Fig. 2B), the H2B density is significantly higher at the nuclear periphery (Fig. 2C) compared to the nucleoplasm (Fig. 2D). In particular, we observed that chromatin-rich (green arrow in Fig. 2D) and chromatic-poor regions distributed throughout the nucleus. This could be due to greater accessibility of the docker to the imager at nuclear periphery, or, as was previously reported, due to inherent inhomogeneity of the H2B distribution^25^. Assuming it’s the latter, we employ a segmentation scheme, for quantitative assessment of chromatin-rich regions, developed by Barth et al.^26^ After segmentation, we observed amorphous structures (see Fig. 2E and its inset) which can be fitted with an elliptic function to unravel the ratio of the minor to major axis. The eccentricity distribution (Fig. 2F) is relatively broad—over values from 0.4 to 1, with peak value ∼0.84. This suggests that most chromatin-rich regions have elongated, fiber-like shape, not circular. The peak values for major and minor axes are ∼98 nm and ∼68 nm respectively (Fig. 2H and 2I). The area distribution (Fig. 2G) fits with log-normal distribution to a mean value of (5.8 ± 0.1) × 10^−3^ *µm*^2^. Taken together, our result strongly suggests that chromatin is spatially distributed in an amorphous structure, shaped like a fiber, with diameters in the sub-100nm range, in excellent agreement with previously reported results^26^.

**Figure 2:**
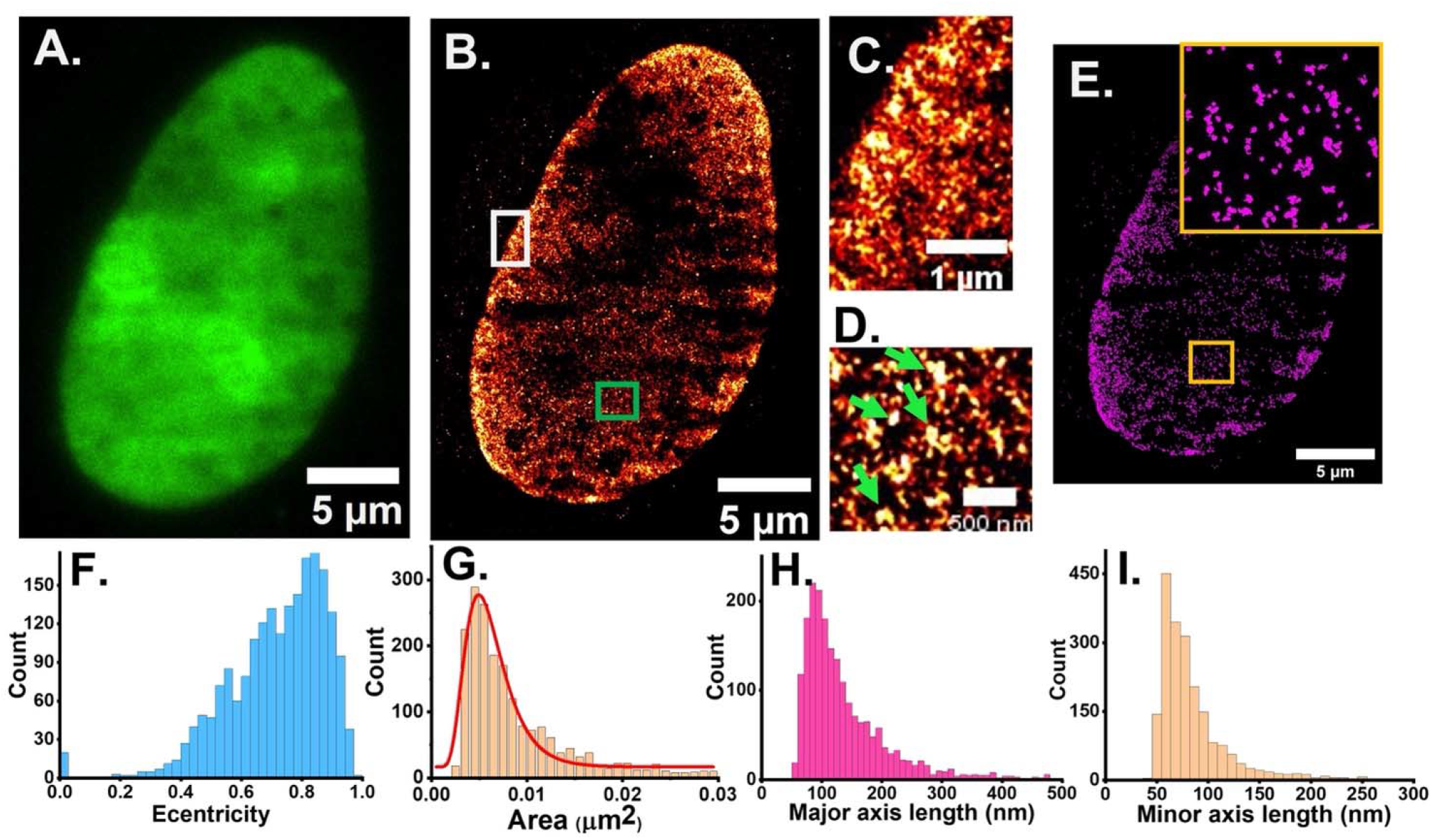
Peptide-PAINT imaging of histone protein H2B via direct conjugation of the E22 docker and transiently binding of a fluorescently labeled imager, LD655-CK19, on fixed HeLa cells. The diffraction-limited image was obtained using GFP, and super-resolution image by peptide-PAINT, using 0.5 nM imager in PBS supplemented with 2% BSA. (A, B): Representative diffraction-limited and super-resolution images of histone protein H2B. (C): Magnified image of perinuclear region, marked as white rectangle. (D): Magnified image of intra-nuclear region, marked as green rectangle in Fig. 2B. (E): Image of H2B-rich regions shown in Fig. 1B after segmentation analysis; inset is magnified region of yellow rectangle. (F, G, H, I): Eccentricity, area (red line: log normal fitted trace); major axis and minor axis distributions of H2B-rich clusters shown in Fig. 2E.

We next attempted to perform nanometric resolution on actin and microtubules, two filamentous cytoskeletal proteins. Following the successful procedure outlined for vimentin and Golgi, we genetically modified the actin-GFP or microtubule-GFP with a E22 docker coil. (The GFP was present to enable a diffraction-limited image). However, the imager did not bind specifically to the docker (data not shown), most likely due to inaccessibility of the docker when in the filament structure of actin or microtubules. Therefore, we co-transfected the plasmid coding for actin-GFP or microtubule-GFP with a separate plasmid coding for the actin or microtubule binding protein appended to the E22 docker. We chose Lifeact, a 17-amino-acid peptide (∼ 1.9 kDa)^27^ for actin, and the microtubule-associated protein-4 (MAP4, ∼43 kDa) for microtubules (Fig. 3A). Notice that this procedure still has the advantage of transfecting a docker into the cell.

**Figure 3:**
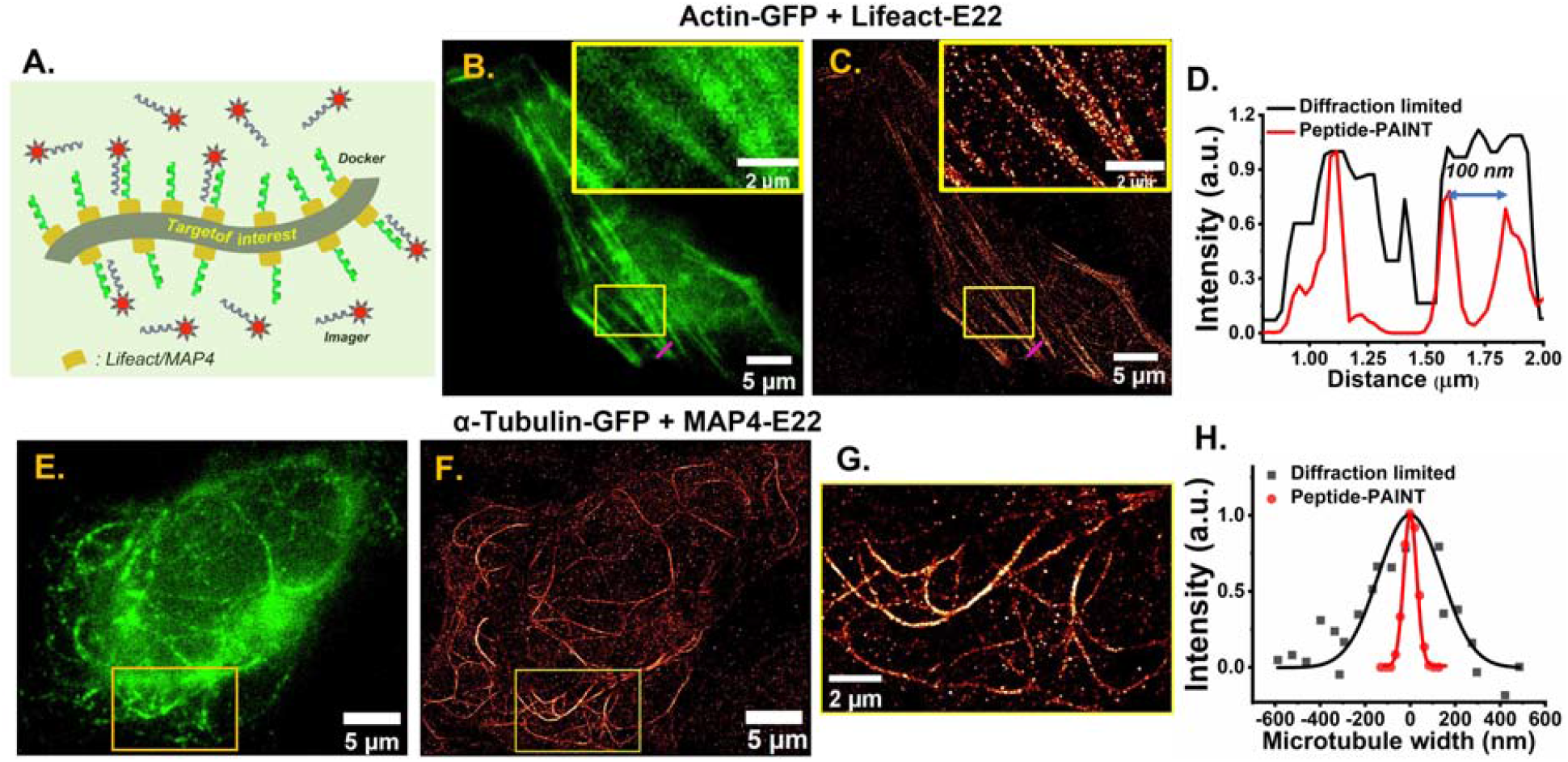
Peptide-PAINT imaging of filament proteins (actin and microtubules) in fixed HeLa cells. In each case, diffraction-limited images were obtained by using GFP fluorescence, and super-resolution images by peptide-PAINT, using 0.5 nM LD655-CK19 in PBS supplemented with 2% BSA as imager coil. (A). Schematic of the experiment. (B-D) For actin imaging, cells were transiently transfected with actin-GFP and Lifeact-E22 plasmids. (B, C) represent diffraction-limited and super-resolution images of actin respectively. (Inset in Figure B, C): Magnification of yellow rectangular region in the respective figures. (D). Comparison of line profile between diffraction-limited (black) and super-resolution (red) images of actin along the magenta lines shown in Figure 2B and 2C. (E-H) For microtubule imaging, cells were transfected with αTubulin-GFP and MAP4-E22 plasmids. (E, F) represent diffraction-limited and super-resolution images of microtubules respectively. (G): Magnification of yellow rectangular region in 2F. (H). Representative traces of microtubule width distributions obtained from diffraction-limited image (solid black rectangle: experimental; black line: Gaussian fitted trace) and super-resolution image (solid red circle: experimental; red line: Gaussian fitted trace).

For actin, both the diffraction-limited (Fig. 3B) and the reconstructed high-resolution (Fig. 3C) images clearly show its filament structures. A negative control (transfected with only an actin-GFP plasmid) shows the actin filament structure only in the diffraction-limited image of the GFP (Fig. S1A), but not in the reconstructed super-resolution image (Fig. S1D). This suggests that the super-resolution images from cells transfected with both actin-GFP *and* Lifeact-E22 plasmids are obtained by specific binding-unbinding events between the docker and the imager. The magnified image of the yellow rectangular box (inset of Fig. 3C) shows a highly resolved structure of actin filaments compared to its corresponding diffraction-limited image (inset of Fig. 3B). This diffraction-limited image is quantified in Fig. 3D, in the region highlighted by the magenta lines in Fig. 3B and 3C, where two filaments are adjacent to each other. In Fig. 3D, we observe two peaks separated by ∼100 nm for the Peptide-PAINT trace while a single broad peak exists for the diffraction-limited trace. (Other results show similar data for other regions.) Hence, our results establishes that Peptide-PAINT can be used to obtain super-resolved structures of actin filaments using a transfected docker on a separate protein.

For microtubules, we had to use an antigen retrieval step, similar to that used with Histone H2B, to create a super-resolution image with Peptide-PAINT. Simply adding the protein-and-docker (tubulin-GFP-E22) and the imager (CK19-LD655) did not yield signal, presumably because of a conformational problem (data not shown). Instead, we transfected tubulin-GFP and MAP4-E22 plasmids and then treated with 1N HCl after fixation and permeabilization (Fig. S6). Fig. 3E shows the diffraction-limited image. This has fewer and discontinuous filament microtubules because at lower pH, irreversible conformational change of GFP decreases its fluorescence^28^ (Fig. 3E). However, in Fig. 3G, the super-resolution image by Peptide-PAINT after antigen retrieval, we observe more resolved and continuous structures. The full-width-half-maximum (FWHM) for the diffraction-limited image is 322 ± 46 nm, while the reconstructed image is 68 ± 9 nm (Figure 3H and S7). This result is very similar to the previously reported STORM study^29^, and shows the improvement in resolution compared to the diffraction-limited image.

Finally, we demonstrate our method for imaging surface proteins in live neurons and cultured HeLa cells to reveal their nanoscale distribution (Fig. 4). Since the imager is impermeable to the cell membrane, we targeted surface proteins such as GluA2, a subunit of an ionotropic glutamate receptor, α-amino-3-hydroxy-5-methyl-4-isoxazolepropionic acid receptor (AMPAR) and neuroligin, a cell adhesion protein on the postsynaptic membrane^30^. GluA2 distribution in the postsynaptic membranes in spines play a key role in synaptic transmission and information processing by the brain^31^. Since it is often present at high density, diffraction-limited imaging cannot resolve its distribution.

**Figure 4:**
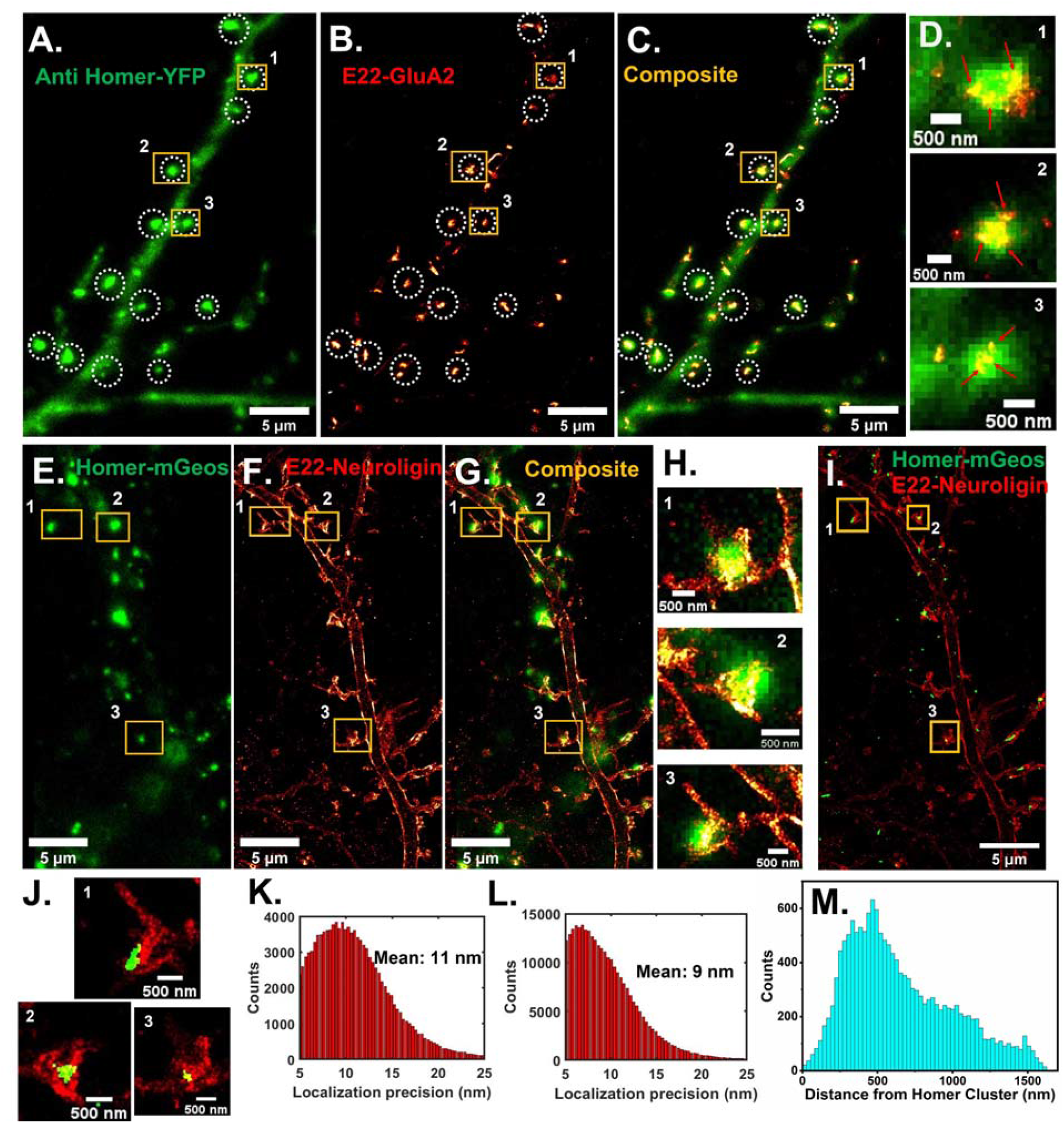
Imaging GluA2-AMPAR and neuroligin in live neurons by peptide-PAINT. DIV14 rat hippocampal neurons were transiently transfected with anti-Homer YFP and E22-GluA2 plasmids for GluA2-AMPAR (A-D, K), and with Homer-mGeos and E22-neuroligin plasmids for neuroligin imaging. (E-J, L, M). (A) Diffraction-limited image of Homer (Green), (B) super-resolution image of GluA2 (red hot), (C) their composite and (D) Zoomed in view of regions 1, 2 and 3 in Figure 4C, showing the colocalization (yellow) respectively. (E) Diffraction-limited image of Homer (Green), (F) super-resolution image of neuroligin (red hot), (G) their composite and (H) Zoomed in view of regions 1,2 and 3 in Figure 4G, marked as yellow rectangles. (I) Composite of super-resolution image of Homer (green) and neuroligin (red). (J) Zoomed in view of regions 1, 2 and 3 in Figure 4I. (K, L) is the localization precision of Fig 4B and 4F respectively. (M) Distribution of neuroligin distances from nearby Homer cluster.

Here, we employ Peptide-PAINT to achieve super-resolution under physiological conditions. To do this, we directly appended the E22-docker to the N-terminal of GluA2, which is exposed to the extracellular side of the cell membrane. We delivered the plasmid to rat hippocampal neurons through transient transfection on day 14 (DIV14). We also transfected another plasmid that encodes an anti-Homer nanobody coupled to a yellow fluorescent protein (YFP)^32^. Homer1 is a post-synaptic protein and the YFP shows the synapse to be fluorescently labeled (at the diffraction-limited resolution) (Fig. 4A, green spots with white-dotted circles). The Peptide-PAINT of GluA2 (Fig. 4B) shows the high-resolution of GluA2. The overlapped images (Fig. 4C, and 4D) shows their co-localizations, indicating specific binding-unbinding between the docker-E22 appended to GluA2 and the imager. The photon counts and localization precision are ∼1325 (Fig. S8A and SI, Table-1) and ∼11 nm (Fig. 4I), respectively. In the magnified images of spines (Fig. 4D), the small clusters of GluA2, indicated by red arrows, are clearly visible, which is due to binding-unbinding events among individual GluA2 or its nanodomains. This establishes that our method can resolve the nanostructures even on live cells. However, we note that the achievable binding frequency for this docker-imager pair is not high enough to acquire sufficient binding-unbinding statistics within the experimental timescale in live cell. Therefore, measuring the number of GluA2 is not possible; furthermore, the bound time for the imager is relatively small (∼220 ms) to measure the diffusion constants. Therefore, designing new pairs of relatively longer binding time (∼0.5-1 sec) may be suitable for single molecule tracking.

As a second example of Peptide-PAINT on a surface protein in live neurons, we chose Neuroligin (NLGN1), a postsynaptic adhesion protein that binds to its presynaptic partner neurexin. We expressed the docker E22 appended to neuroligin’s N-terminus in neurons through transient transfection of a plasmid encoding their sequences. Another plasmid, Homer-mGeos, was used as a transfection- and synaptic-marker. As expected, we observed the exchange of the imager in cells positive for mGeos and co-localization between Homer clusters (which are diffraction-limited) and neuroligin (Fig. 4G and 4H). This ensures the specific binding-unbinding of the imager to the docker (Fig. 4A-D). Since mGeos is a super-resolution GFP-like marker (unlike GFP), we can obtain the super-resolution image of Homer clusters (Fig. 4I and J, S9) and to determine the distance between Homer and neuroligin (Fig. 4M). Neuroligin is broadly distributed with the most probable distance ∼410 nm from the neighboring Homer cluster, and ∼82% of neuroligin at the synapse is localized within 1µm distance from the Homer cluster (Fig. S10). The photon counts and localization precision are ∼1453 (Fig. S8B) and ∼9 nm (Fig. 4L) respectively. We also tested neuroligin imaging in live HeLa cells and obtained similar result (Fig. S11). Taken together, we achieve robust Peptide-PAINT images on the surface of live neurons and HeLa cells using our method.

## CONCLUSION

In conclusion, we have introduced a novel method for Peptide-PAINT for achieving nanometric resolution based on E22-CK19 coiled-coil pair and we have extended Peptide-PAINT to look at live neurons and mammalian cells under physiological conditions. We demonstrate that the internally expressed docker to various targets via transient transfection, enables Peptide-PAINT super-resolution imaging of different organelles including vimentin, Golgi body, H2B, actin, and microtubules in fixed HeLa cells, and surface proteins, namely GluA2 and neuroligin in *live* neurons. To our knowledge, this is the first demonstration of Peptide-PAINT imaging in live mammalian cells. In some cases with fixed cells, it was necessary to do antigen retrieval and in other cases it was necessary to use two plasmids: one containing the original targeted protein and the other containing a target binding protein. With the super-resolution image by Peptide-PAINT, we uncovered key nanoscale structural details of different biological targets. Specifically, we found a ∼60 nm periodicity in the unmutated vimentin filaments inside cells, supporting longer vimentin filaments formation by longitudinal assembly of ULFs. We also found sub-100 nm fiber-shaped amorphous structures of chromatin rich regions in the nucleus of HeLa cells. Finally, we measured in live cells under physiological buffer conditions, GluA2 and neuroligin distributions at the synapses in neurons. We argue that our method is easy-to-use compared to conventional DNA-PAINT owing to elimination of time-consuming multiple steps for the docker labeling.

## ASSOCIATED CONTENT

### Supporting information

Additional detailed descriptions of materials, experimental methods, and figures, plasmid sequences

## ACKNOWLEDGEMENTS

This work is supported by NIH Grant R01 GM132392 and by the NSF Physics Frontiers Center (PFC) grant PHY-1430124.

## REFERENCES

1. Rust, M. J., Bates, M. & Zhuang, X. Sub-diffraction-limit imaging by stochastic optical reconstruction microscopy (STORM). Nat. Methods 3, 793–795 (2006).

2. Betzig, E. et al. Imaging intracellular fluorescent proteins at nanometer resolution. Science (80-.). 313, 1642–1645 (2006).

3. Yildiz, A. et al. Myosin V Walks Hand-Over-Hand: Single Fluorophore Imaging with 1.5-nm Localization. Science (80-.). 300, 2061–2065 (2003).

4. Strauss, S. et al. Modified aptamers enable quantitative sub-10-nm cellular DNA-PAINT imaging. Nat. Methods 15, 685–688 (2018).

5. Hess, S. T., Girirajan, T. P. K. & Mason, M. D. Ultra-high resolution imaging by fluorescence photoactivation localization microscopy. Biophys. J. 91, 4258–4272 (2006).

6. Sharonov, A. & Hochstrasser, R. M. Wide-field subdiffraction imaging by accumulated binding of diffusing probes. Proc. Natl. Acad. Sci. U. S. A. 103, 18911–18916 (2006).

7. Hell, S. W. Far-field optical nanoscopy. Science (80-.). 316, 1153–1159 (2007).

8. Strauss, S. & Jungmann, R. Up to 100-fold speed-up and multiplexing in optimized DNA-PAINT. Nat. Methods 17, 789–791 (2020).

9. Schueder, F. et al. An order of magnitude faster DNA-PAINT imaging by optimized sequence design and buffer conditions. Nat. Methods 16, 1101–1104 (2019).

10. Eklund, A. S., Ganji, M., Gavins, G., Seitz, O. & Jungmann, R. Peptide-PAINT Super-Resolution Imaging Using Transient Coiled Coil Interactions. Nano Lett. 20, 6732–6737 (2020).

11. Jungmann, R. et al. Multiplexed 3D cellular super-resolution imaging with DNA-PAINT and Exchange-PAINT. Nat. Methods 11, 313–8 (2014).

12. Unterauer, E. M. & Jungmann, R. Quantitative Imaging With DNA-PAINT for Applications in Synaptic Neuroscience. Front. Synaptic Neurosci. 13, 1–8 (2022).

13. Jungmann, R. et al. Quantitative super-resolution imaging with qPAINT. Nat. Methods 13, 439–442 (2016).

14. Fischer, L. S. et al. Quantitative single-protein imaging reveals molecular complex formation of integrin, talin, and kindlin during cell adhesion. Nat. Commun. 12, (2021).

15. Oi, C. et al. LIVE-PAINT allows super-resolution microscopy inside living cells using reversible peptide-protein interactions. Nat. Commun. Biol. 3:458, (2020).

16. Fischer, L. S., Schlichthaerle, T., Chrostek-Grashoff, A. & Grashoff, C. Peptide-PAINT Enables Investigation of Endogenous Talin with Molecular Scale Resolution in Cells and Tissues. ChemBioChem 22, 2872–2879 (2021).

17. Perfilov, M. M., Gavrikov, A. S., Lukyanov, K. A. & Mishin, A. S. Transient fluorescence labeling: Low affinity—high benefits. Int. J. Mol. Sci. 22, (2021).

18. Oi, C. S. G. J. M. PAINT using proteins: A new brush for super-resolution artists. Protein Sci. 29, 2142–2149 (2020).

19. Tas, R. P., Albertazzi, L. & Voets, I. K. Small Peptide-Protein Interaction Pair for Genetically Encoded, Fixation Compatible Peptide-PAINT. Nano Lett. 21, 9509–9516 (2021).

20. Volk, L., Chiu, S. L., Sharma, K. & Huganir, R. L. Glutamate Synapses in Human Cognitive Disorders. Annu. Rev. Neurosci. 38, 127–149 (2015).

21. Schnitzbauer, J., Strauss, M. T., Schlichthaerle, T., Schueder, F. & Jungmann, R. Super-resolution microscopy with DNA-PAINT. Nat. Protoc. 12, 1198–1228 (2017).

22. Ovesný, M., Křížek, P., Borkovec, J., Švindrych, Z. & Hagen, G. M. ThunderSTORM: A comprehensive ImageJ plug-in for PALM and STORM data analysis and super-resolution imaging. Bioinformatics 30, 2389–2390 (2014).

23. Peters, R., Griffié, J., Burn, G. L., Williamson, D. J. & Owen, D. M. Quantitative fibre analysis of single-molecule localization microscopy data. Sci. Rep. 8, 1–8 (2018).

24. Vicente, F. N. et al. Molecular organization and mechanics of single vimentin filaments revealed by super-resolution imaging. Sci. Adv. 8, 1–16 (2022).

25. Nozaki, T. et al. Dynamic Organization of Chromatin Domains Revealed by Super-Resolution Live-Cell Imaging. Mol. Cell 67, 282-293.e7 (2017).

26. Barth, R., Bystricky, K. & Shaban, H. A. Coupling chromatin structure and dynamics by live super-resolution imaging. Sci. Adv. 6, (2020).

27. Riedl, J. et al. Lifeact: A versatile marker to visualize F-actin. Nat. Methods 5, 605–607 (2008).

28. Kneen, M., Farinas, J., Li, Y. & Verkman, A. S. Green fluorescent protein as a noninvasive intracellular pH indicator. Biophys. J. 74, 1591–1599 (1998).

29. Bates, M., Huang, B., Dempsey, G. T. & Zhuang, X. Multicolor super-resolution imaging with photo-switchable fluorescent probes. Science (80-.). 317, 1749–1753 (2007).

30. Song, J. Y., Ichtchenko, K., Südhof, T. C. & Brose, N. Neuroligin 1 is a postsynaptic cell-adhesion molecule of excitatory synapses. Proc. Natl. Acad. Sci. U. S. A. 96, 1100–1105 (1999).

31. Chater, T. E. et al. The role of AMPA receptors in postsynaptic mechanisms of synaptic plasticity. Front. Cell. Neurosci. 8, 1–14 (2014).

32. Dong, J. X. et al. A toolbox of nanobodies developed and validated for use as intrabodies and nanoscale immunolabels in mammalian brain neurons. Elife 8, 1–25 (2019).

33. Tas, R. P., Albertazzi, L. & Voets, I. K. Small Peptide–Protein Interaction Pair for Genetically Encoded, Fixation Compatible Peptide-PAINT. Nano Lett. (2021) doi:10.1021/acs.nanolett.1c02895.

34. Hayashi, T., Lewis, A., Hayashi, E., Betenbaugh, M. J. & Su, T. P. Antigen retrieval to improve the immunocytochemistry detection of sigma-1 receptors and ER chaperones. Histochem. Cell Biol. 135, 627–637 (2011).

35. Tang, X., Falls, D. L., Li, X., Lane, T. & Luskin, M. B. Antigen-retrieval procedure for bromodeoxyuridine immunolabeling with concurrent labeling of nuclear DNA and antigens damaged by HCl pretreatment. J. Neurosci. 27, 5837–5844 (2007).

36. Laporte, M. H., Klena, N., Hamel, V. & Guichard, P. Visualizing the native cellular organization by coupling cryofixation with expansion microscopy (Cryo-ExM). Nat. Methods 19, 216–222 (2022).

37. Manley, S. et al. High-density mapping of single-molecule trajectories with photoactivated localization microscopy. Nat. Methods 5, 155–157 (2008).

