## Supplementary material for "Peptide-PAINT using a transfected-docker enables live- and fixed-cell super-resolution imaging": Supporting information.docx

**Methods**

**Peptide-dye labelling**

Imager peptide (K19) with an additional cysteine in the N-terminal was named as CK19. This was purchased from GenScript, USA. It was dissolved in PBS at pH 7.4 with concentration 1 mg/ml. 100 µL of 40 µM peptide solution was incubated with 250 µM tris(2-carboxyethyl) phosphine (TCEP) for 30 min at room temperature. Afterwards, 5-fold excess of LD655-maleimide dye (Lumidyne Technologies, USA) dissolved in 4 µL DMSO was added to peptide solution and incubated at 4C for overnight. LD655 labeled CK19 was separated from free dye using HPLC. The solvent was evaporated, and then the labeled peptide was dissolved in PBS and used for imaging experiments.

**Plasmid construction**

H2B-GFP-E22: H2B-Halo Tag was a gift from Kai Johnsson (Addgene plasmid # 169329). The sequence except Halo tag was copied from this plasmid, and GFP sequence was copied from Vimentin-EGFP plasmid (Addgene plasmid # 56439) using PCR. Then, both were assembled by HiFi assembly to make H2B-GFP plasmid. Afterwards, the docker E22 sequence was inserted at the C-terminal of GFP by Q5 site-directed mutagenesis.

Golgi7-GFP-E22: Golgi7-dsRed plasmid was a gift from Michael Davidson (Addgene plasmid #55832). Golgi7 sequence was copied from it and GFP-E22 sequence with backbone sequence was copied from H2B-GFP-E22 plasmid using PCR. Both were assembled by HiFi assembly to make Golgi7-GFP-E22 plasmid.

Vimentin-GFP-E22: Vimentin-EGFP plasmid was a gift from Michael Davidson (Addgene plasmid # 56439). The docker E22 sequence was inserted at the C-terminal of GFP by Q5 site-directed mutagenesis.

Lifeact-E22: mTagBFP-Lifeact-7 plasmid was a gift from Michael Davidson (Addgene plasmid # 54496). mTagBFP sequence was deleted and E22 sequence was inserted by Q5 site-directed mutagenesis in two sequential steps.

Actin-GFP: mEos2-Actin-N-10 was a gift from Michael Davidson (Addgene plasmid # 57341). mEos2 sequence was deleted by Q5 site-directed mutagenesis. Afterwards, GFP sequence, copied from H2B-GFP-E22 plasmid, was inserted at the C-terminal of actin using HiFi assembly.

a-tubulin-GFP: mScarlet-tubulin plasmid was a gift from Dorus Gadella (Addgene plasmid #85047). mScarlet sequence was deleted by Q5 site-directed mutagenesis. GFP sequence was copied from H2B-GFP-E22 plasmid. Then, both were assembled using HiFi assembly.

MAP4-E22: MAP4-iFAST plasmid was a gift from Arnaud Gautier (Addgene plasmid #130820) iFAST sequence was replaced by the docker E22 sequence using Q5 site-directed mutagenesis.

E22-GluA2: AP-GluA2 plasmid was a gift from our collaborator (Prof. William Green, University of Chicago, USA). The AP sequence was substituted by the docker E22 sequence using Q5 site-directed mutagenesis.

E22-Neuroligin: To construct the E22-Neuroligin, we substituted the HA epitope of HA-Neuroligin (Addgene Plasmid #15260) by the docker-E22 sequence using Q5 site-directed mutagenesis.

aHomer-YFP: HC20 pEYFPN1 anti-Homer1 nanobody was a gift from James Trimmer (Addgene plasmid # 135220). YFP sequence was copied from pCAG-YFP, a gift from Connie Cepko (Addgene plasmid #11180;) and then inserted into HC20 pEYFPN1 anti-Homer1 nanobody plasmid by HiFi assembly.

mGeos- Homer: mGeos-M-Homer1-C-18 was a gift from Michael Davidson (Addgene plasmid # 57528).

**Cell culture and fixation for imaging**

HeLa cells were cultured to 70% confluency on 35 mm MatTec dishes in Minimal Essential Media (MEM). Cells were transiently transfected for 2 hours with 1μg of DNA plasmids (Vimentin-GFP-E22/Actin-GFP+Lifeact-E22/αTubulin-GFP+MAP4-E22/H2B-GFP-E22/Golgi-GFP-E22) followed by incubation at 37C for 16-20 hrs. Cells were washed two times with PBS, fixed with 4% PFA for 10 min, and then permeabilized with 0.25% Triton X-100 for 10 min. However, only the MAP4-E22 and αTubulin-GFP transfected cells were fixed with 4% PFA in presence of 0.1% Triton X-100 for 10 min, and then permeabilized with 0.25% Triton X-100 for 10 min. After Triton X-100 wash, cells were treated with 0.5, 1 and 2N HCl for 20 min separately, and for H2B-GFP-E22 transfected cells, with 1% SDS for 10 min to overcome fixative artefacts. Afterwards, cells were treated with the blocking solution (2% BSA+5% sucrose+10% FBS) for 3 hrs followed by washing with PBS three times. Then, imager solution (0.5 nM LD655-CK19 supplemented with 2% BSA) was added to it and the sample was subjected to imaging. Imaging parameters namely total frame numbers, laser power and exposure time have been mentioned in Table-1.

**Live cell Peptide-PAINT imaging of neuroligin in HeLa cells**

HeLa cells were cultured to 70% confluency on 35 mm MatTec dishes in MEM media. Cells were transfected for 4 hours with E22-NLGN and LifeAct-BFP using Lipofectamine 2000. Then, the media was replaced with fresh MEM media. We incubated overnight and imaged the next day. Cells were illuminated in 500 pM LD6550-CK19 imager in HiLo mode^1^ with low 405 laser power for LifeAct-BFP and 640 nm laser (74 W/cm^2^) for 5000 frames of 50 ms exposure time.

**Peptide-PAINT imaging in live neurons**

To prepare primary hippocampal cultures from E18 rats, hippocampal tissues were dissociated in 3 mg/mL protease and plated on 25 mm coverslips coated with 1 mg/mL poly-l-lysine and laminin. Neurons were cultured at 37˚C with 5% CO_2_ in neurobasal media (ThermoFisher, 21103049) with B-27 supplement (GIBCO), 2 mM GlutaMAX (GIBCO) and 50 unit/mL penicillin and 50 unit/ml streptomycin. On 14 days in vitro (DIV), neurons were co-transfected with anti-Homer-YFP (1 mg/coverslip) and E22-GluA2 (1 mg/coverslip) plasmids for GluA2 imaging, and Homer-mGeos and E22-neuroligin plasmids for neuroligin imaging by using Lipofectamine 2000 transfection reagent. After ~48 hrs of transfection, the coverslips were transferred to warm imaging buffer (HBSS supplemented with 10 mM Hepes, 1 mM MgCl_2_, 1.2 mM CaCl_2_ and 2 mM D-glucose) in presence of 5% casein for 5 min incubation and mounted onto an imaging dish (Warner RC-40LP). Neurons are then washed with the imaging buffer, and the imager solution having 500 pM LD655-CK19 was added to it. A diffraction-limited image of Homer clusters has been recorded by exciting YFP with 488 nm laser and collecting fluorescence in the range of 510-560 nm. Then, Peptide-PAINT imaging was done exciting with 640 laser and collecting fluorescence in the range of 660-700 nm with 100 ms exposure time. Detected individual binding event was localized with high precision and employed to reconstruct GluA2 super-resolution image. For neuroligin imaging only, mGeos was activated with 405 nm laser and imaged with 488 nm excitation, and Peptide-PAINT was done using 640 nm laser excitation.

**Microscope set up**

Imaging was carried out with a Nikon TI Eclipse microscope with a Nikon APO 100X oil immersion objective (N.A. 1.49) and an additional 1.5x magnifying lens and with a Perfect Focus System for Z-stabilization. An Agilent laser system MLC400B with four fiber-coupled lasers (405 nm, 488 nm, 561 nm, and 640 nm) was used for illumination. Elements software from Nikon was used for data acquisition. A back illuminated EMCCD (Andor DU897) was used for recording. x-y-z position has been controlled by a motorized stage from ASI with a Piezo top plate (ASI PZ-2000FT). A quad-band dichroic (Chroma, ZT405-488-561-640RPC) and band-pass emission filter 525/50, 780/40 were used for fluorescence imaging.

**Image analysis**

Raw fluorescence data was subjected to super-resolution reconstruction using the ‘Picasso’ software package^2^ as well as ThunderSTORM plugin in Fiji^3^. During cell imaging, bright spot, easily observable in the field of view of transmission image, was used as fiducials for drift correction. The drift correction was performed using home-written python code.

In the super-resolution image of vimentin obtained by Peptide-PAINT, the intensity profile of individual vimentin filaments along the filament axis was obtained from the low-density regions using Fiji, and then fitted with multiple peaks Gaussian function in OriginPro2019.

In the super-resolved image of histone protein H2B obtained by Peptide-PAINT, binary threshold is applied to separate high density amorphous structures. Afterwards, the amorphous structures are identified using watershed segmentation algorithm and fitted with elliptic function to unravel the structural parameters. The complete analysis was performed using Fiji^4^.

The intensity profile across the microtubule for both diffraction-limited and super-resolution image by Peptide-PAINT was fitted with Gaussian function, and full width at half maxima (FWHM) was defined as the microtubule width.

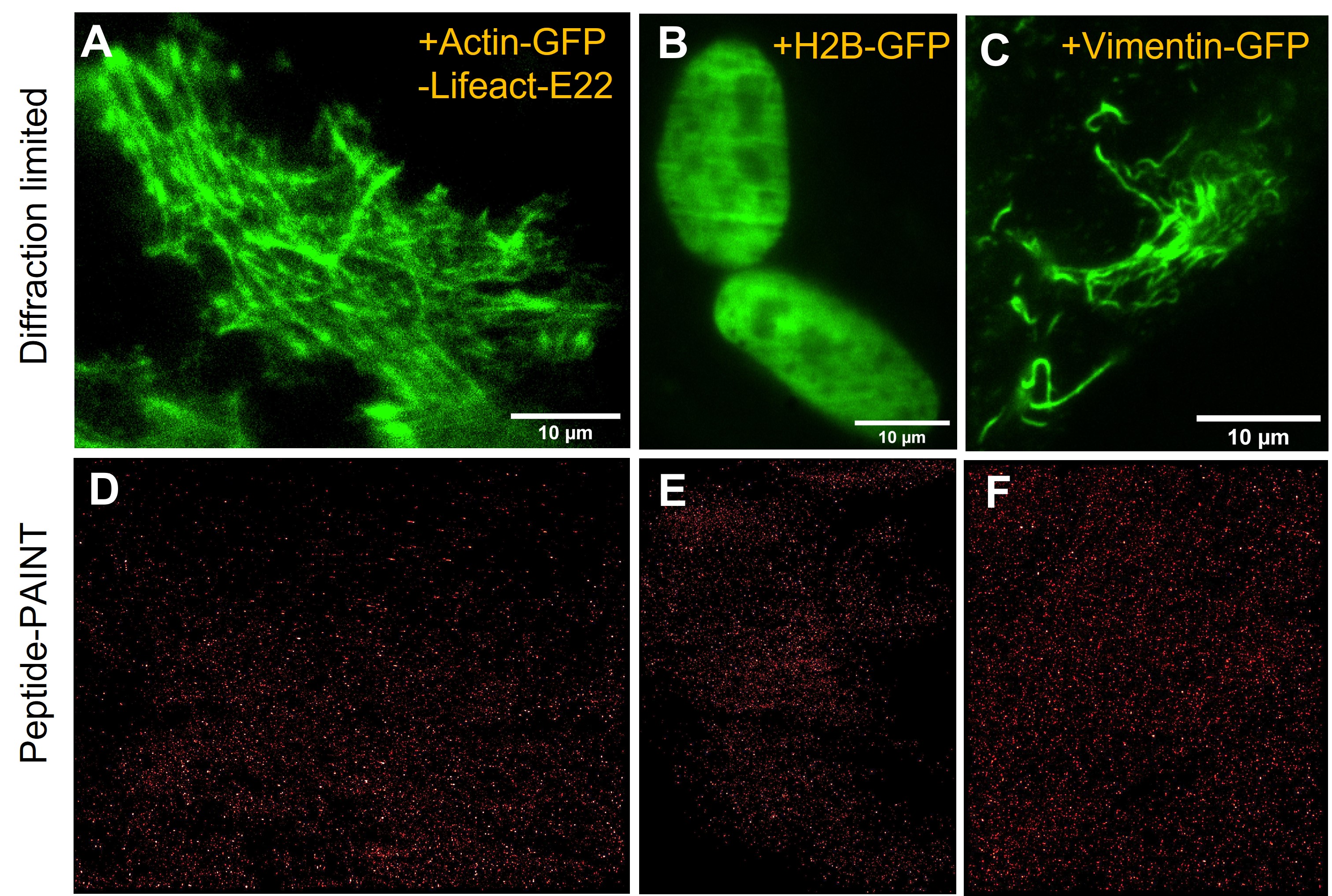

**Fig. S1: Negative control for imaging actin, H2B and vimentin by Peptide-PAINT.** A, B, C: Diffraction limited images for Actin-GFP, H2B-GFP and Vimentin-GFP transfected cells; D, E, F: reconstructed images from Peptide-PAINT for Actin-GFP, H2B-GFP and Vimentin-GFP transfected cells (Scale bar is same for diffraction-limited and the corresponding Peptide-PAINT images.)

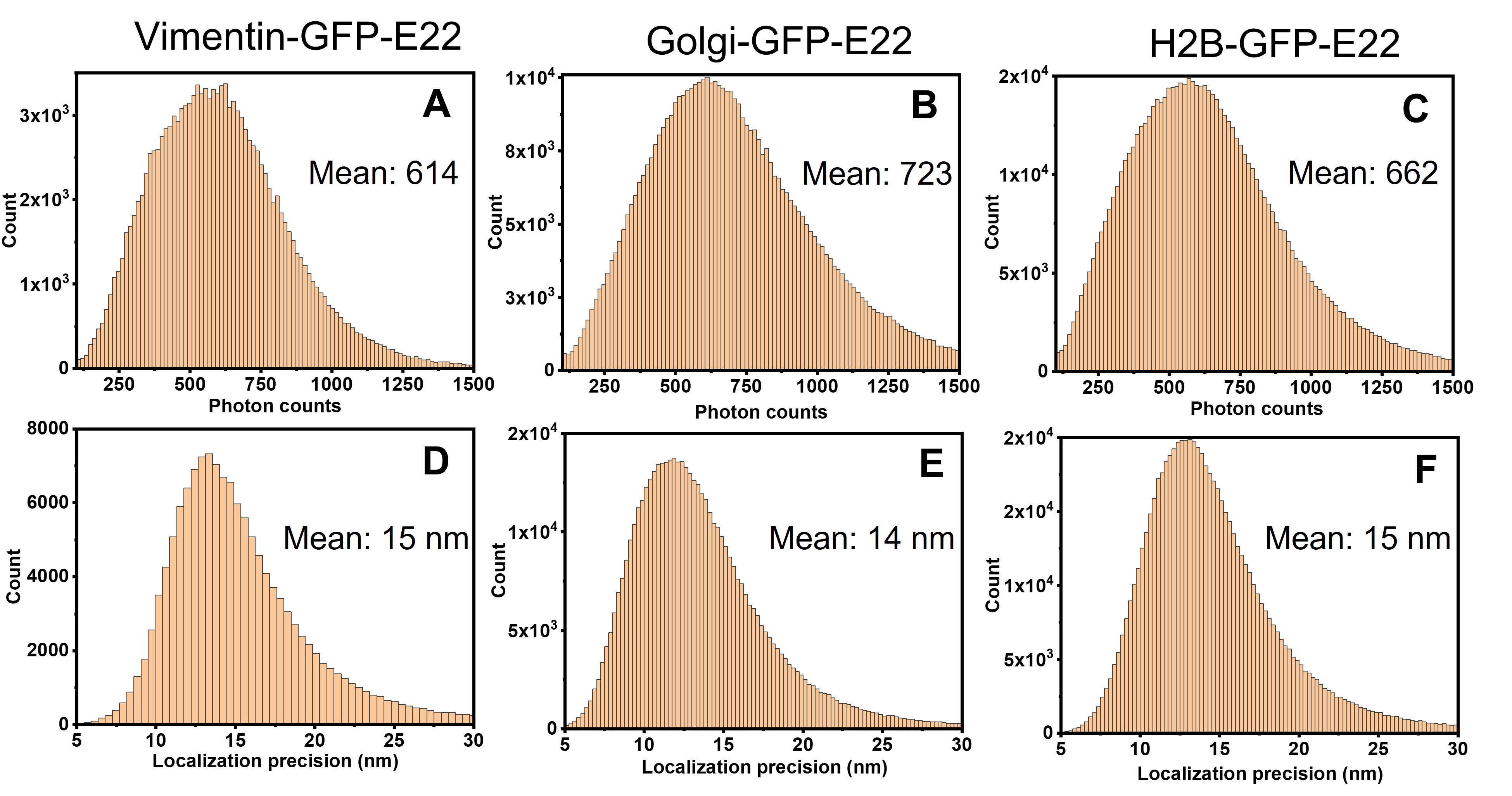

**Fig. S2: Photon counts and localization precision histograms for Peptide-PAINT imaging.** Vimentin-GFP-E22 (A, D), Golgi-GFP-E22 (B, E) and H2B-GFP-E22 (C, F) transfected fixed HeLa cells (Analyzed by Picasso).

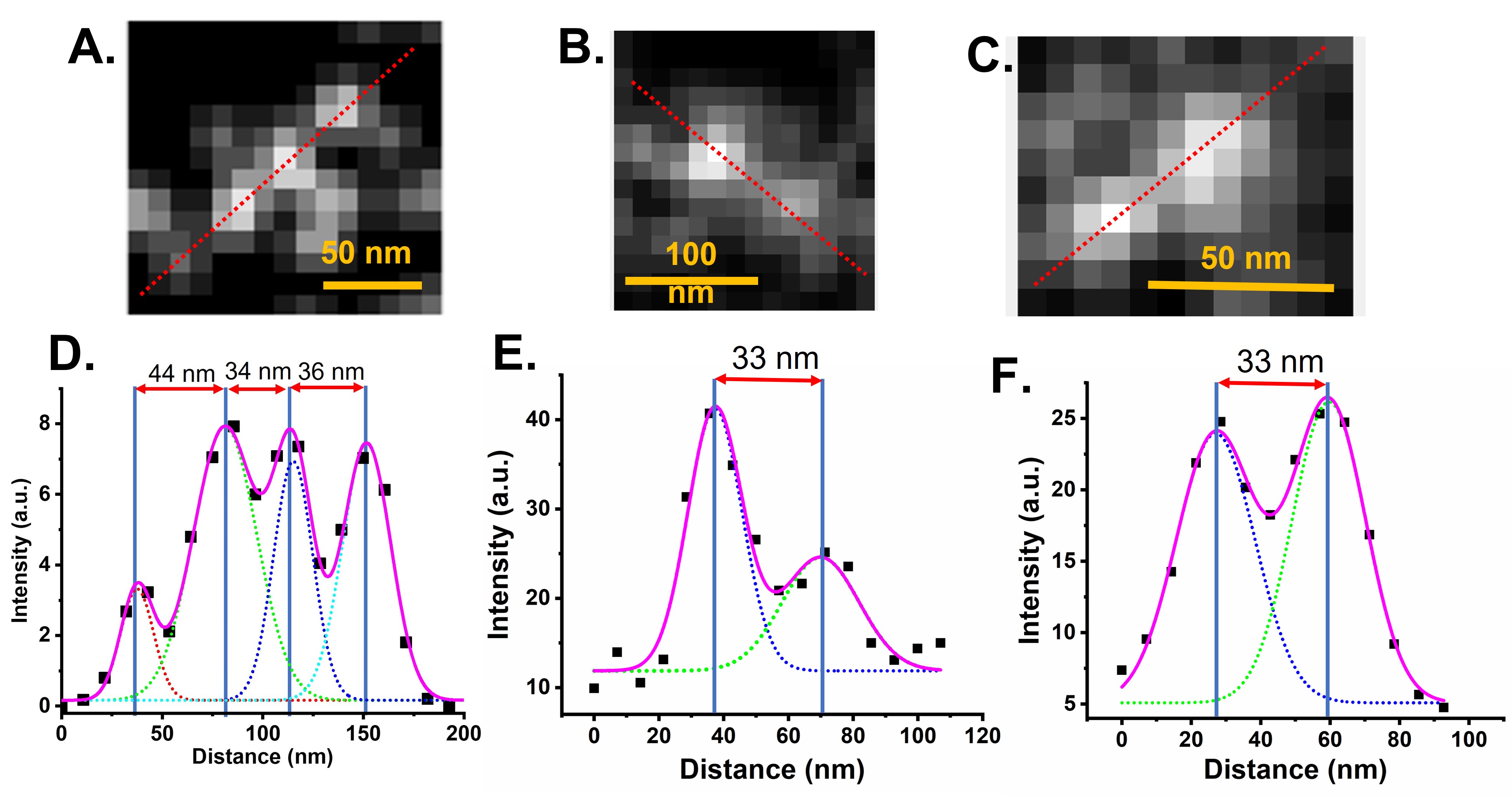

**Fig. S3: Peptide-PAINT imaging of vimentin.** (A, B, C) Representative magnified images single vimentin filament. (D, E, F) Intensity profile along the red-dotted line in Fig. 1A, 1B and 1C respectively.

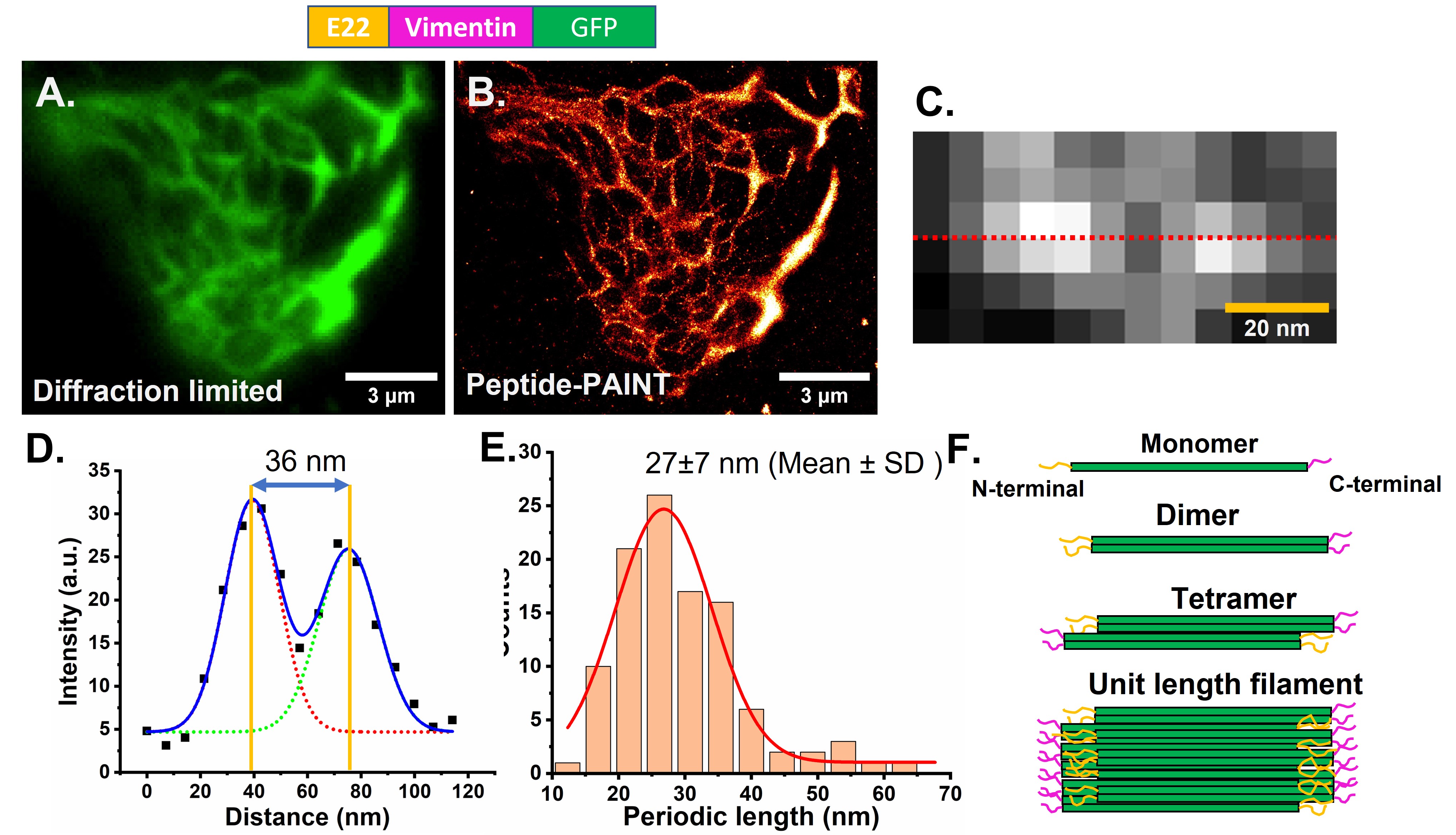

**Fig. S4:** **Peptide-PAINT and Diffraction-limited imaging of vimentin on fixed HeLa cells.** Cells were transfected with E22-Vimentin-GFP plasmid. Diffraction limited images were obtained by using GFP fluorescence, and super-resolution images by Peptide-PAINT using 0.5 nM LD655-CK19 in PBS supplemented with 2% BSA as imager. (A, B) Representative diffraction-limited and super-resolution images of vimentin. (C) Representative magnified image of representing single vimentin filament. (D) Intensity profile along the red-dotted line in Fig. 1C. Intensity profile was fitted with two Gaussian functions to obtain the distance between the centers of two neighboring spots. (E) Periodic length distribution of vimentin filament. (F) Cartoon of vimentin unit length filament formation.

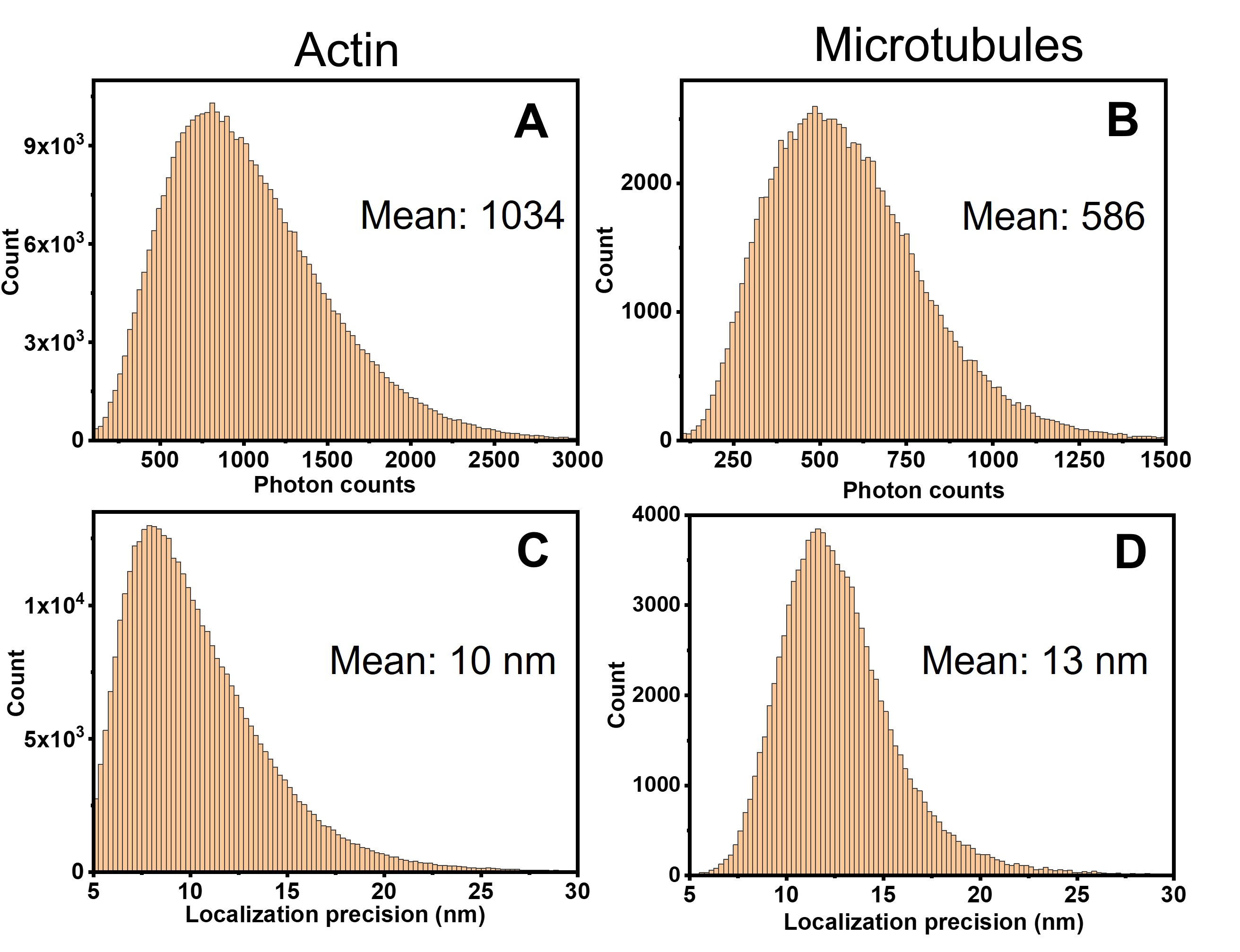

**Fig. S5: Photon counts and localization precision histograms for Peptide-PAINT imaging.** actin-GFP+Lifeact-E22 (A, C), and α-tubulin-GFP+MAP4-E22 (B, D) transfected fixed HeLa cells (Analyzed by Picasso).

**Fig. S5:** Microtubule width (FWHM obtained by Gaussian fitting of intensity profile across the microtubule filament) distribution in HeLa cells transfected with MAP4-E22 and α-tubulin-GFP plasmids and then treated with 1N HCl for antigen retrieval. (A) Diffraction-limited, (B) Super-resolution by Peptide-PAINT.

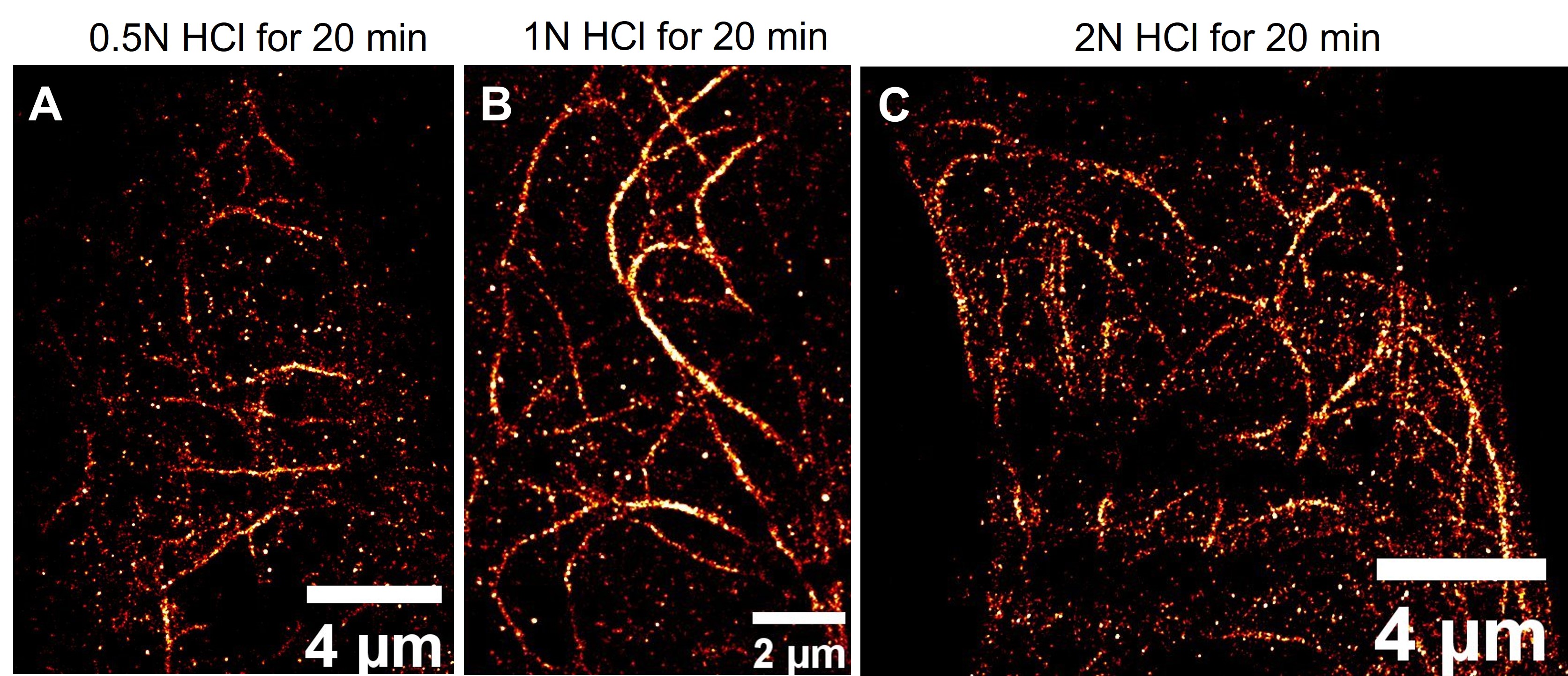

**Fig. S6: Microtubule imaging by Peptide-PAINT under different antigen-retrieval conditions.** In each case, cells were transiently transfected with MAP4-E22 and αTubulin-GFP plasmid. After ~20 hrs of transfection, cells were fixed with 4% PFA and 0.1% Triton X-100 for 10 min followed by permeabilization with 0.25% triton-X100 for 10 min. Then, treated with 0.5 N HCl (A), 1N HCl (B), 2N HCl (C) for 20 min. Cells were incubated with the blocking solution (see Methods for details) and then subjected to Peptide-PAINT imaging in presence of 0.5 nM LD655-CK19 supplemented with 2% BSA.

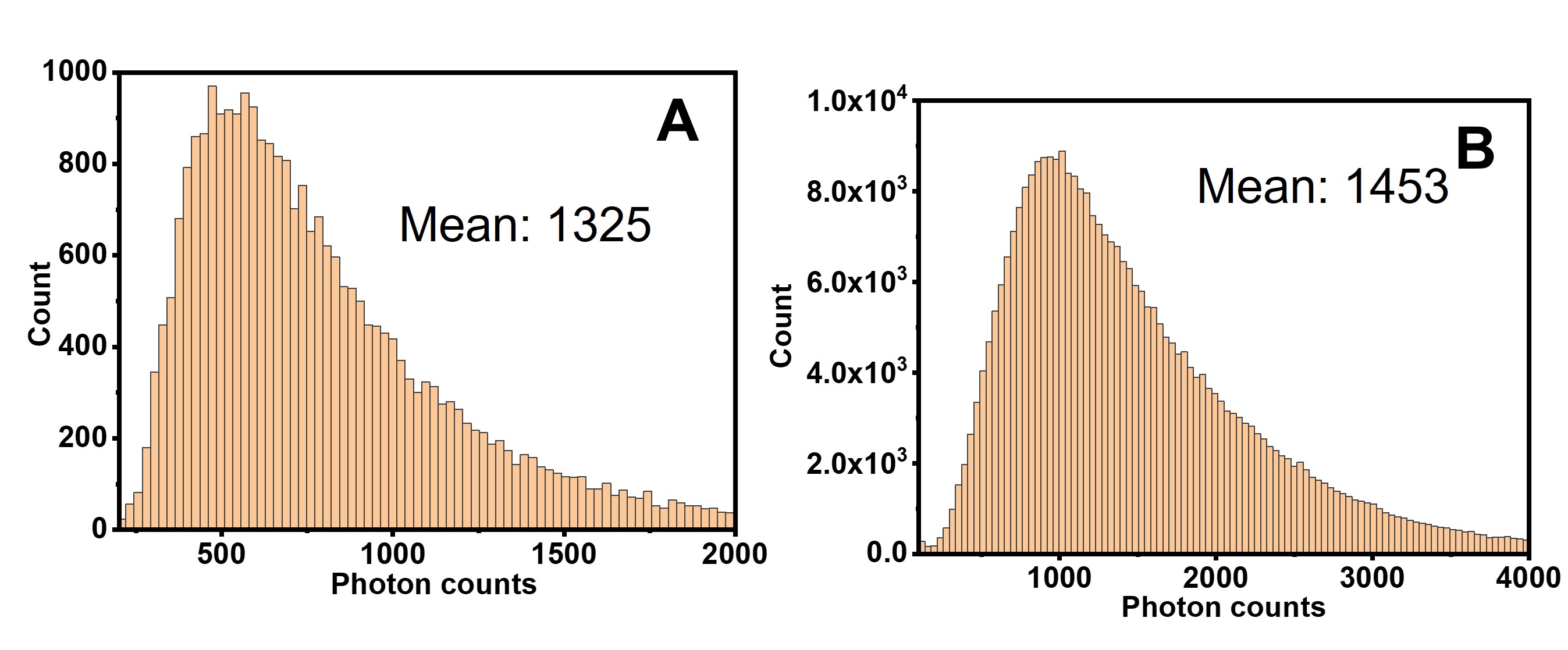

**Fig. S8: Photon counts histograms for Peptide-PAINT imaging.** (A) Anti-Homer-YFP+E22-GluA2, and (B) Homer-mGeos+E22-Neuroligin transfected live neurons (Analyzed by Picasso).

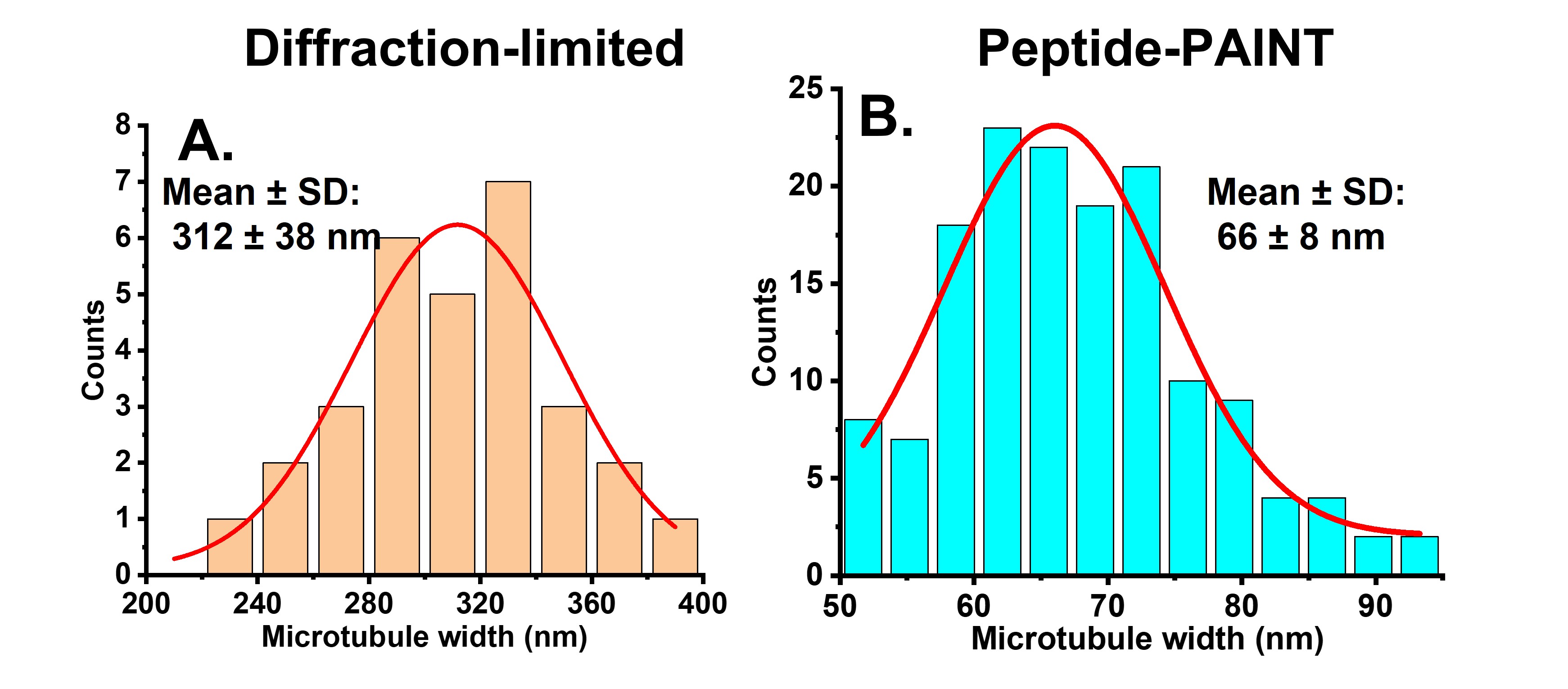

**Fig. S7:** Microtubule width (FWHM obtained by Gaussian fitting of intensity profile across the microtubule filament) distribution in HeLa cells transfected with MAP4-E22 and α-tubulin-GFP plasmids and then treated with 1N HCl for antigen retrieval. (A) Diffraction-limited, (B) Super-resolution by Peptide-PAINT.

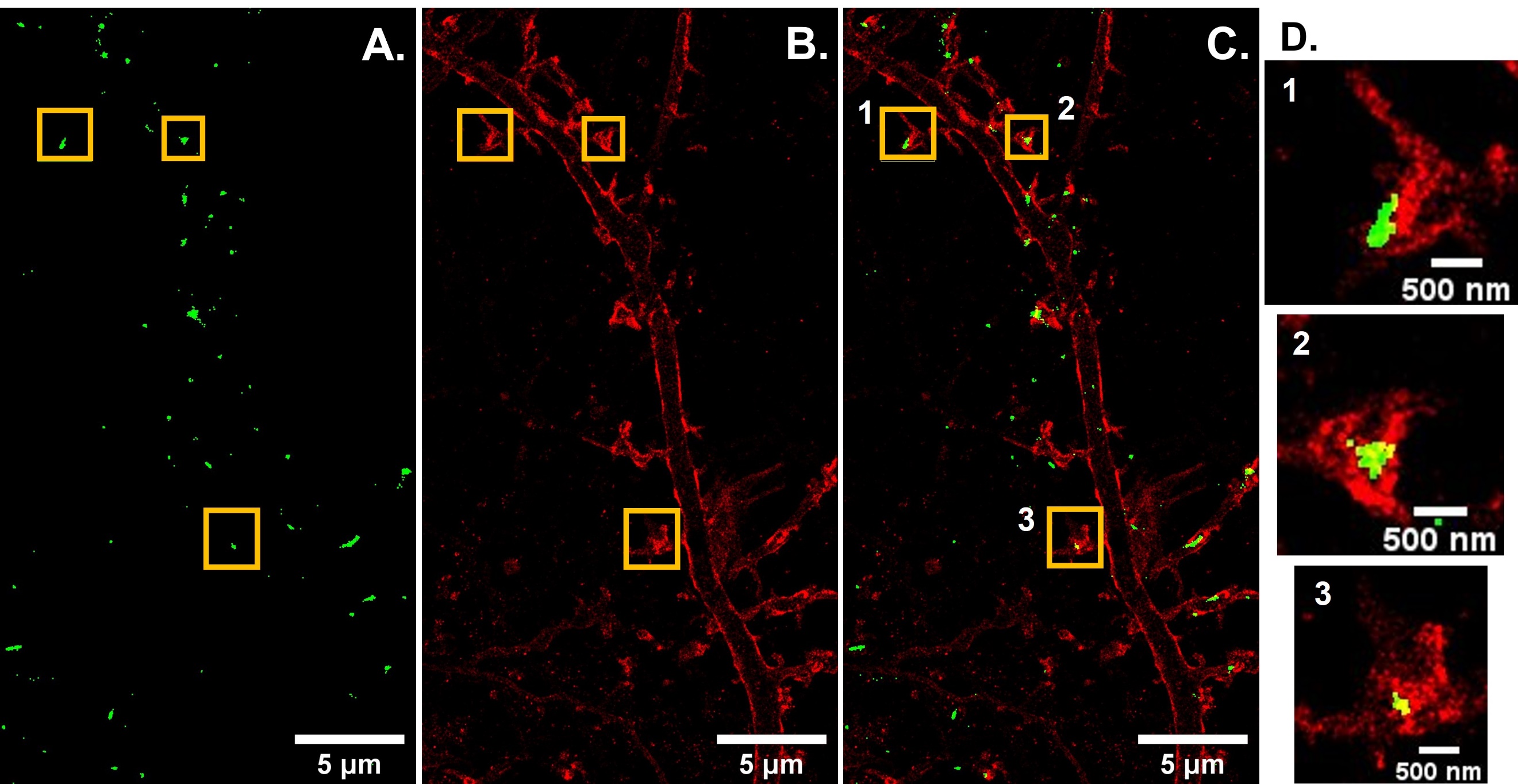

**Fig. S9:** **Imaging Homer by photoactivated localization microscopy (PALM) and** **neuroligin in live neurons by peptide-PAINT.** DIV14 rat hippocampal neurons were transiently transfected with Homer-mGeos and E22-neuroligin plasmids**.** (A) Super-resolution image of Homer by PALM (Green), (B) super-resolution image of neuroligin by Peptide-PAINT (red), (C) their composite and (D) Magnified view of regions 1,2 and 3 in Figure S7C, marked as yellow rectangles.

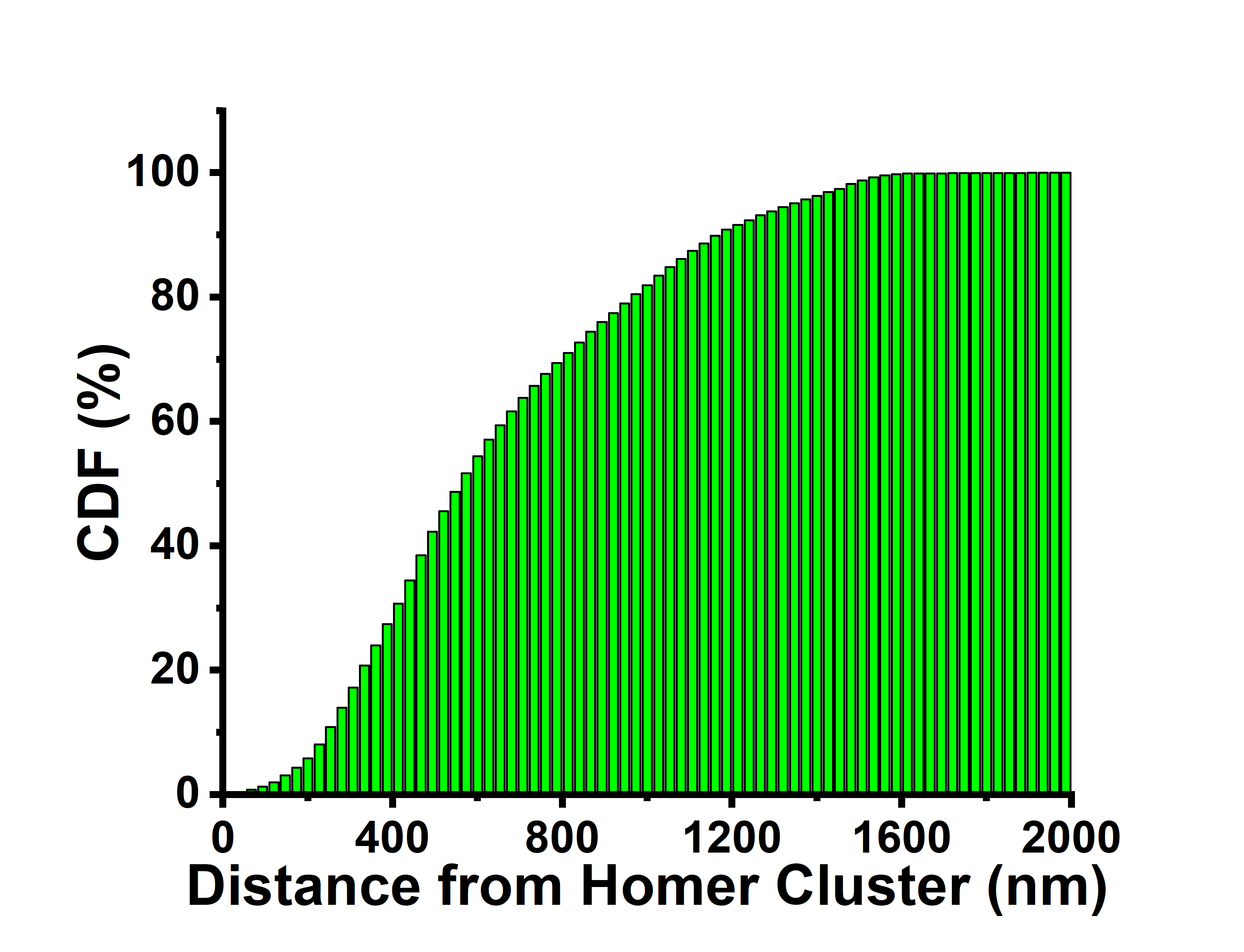

**Fig. S10: Cumulative distribution function (CDF) of distances between neuroligin and Homer clusters.** This has been calculated from Fig. S9.

**Table-1**

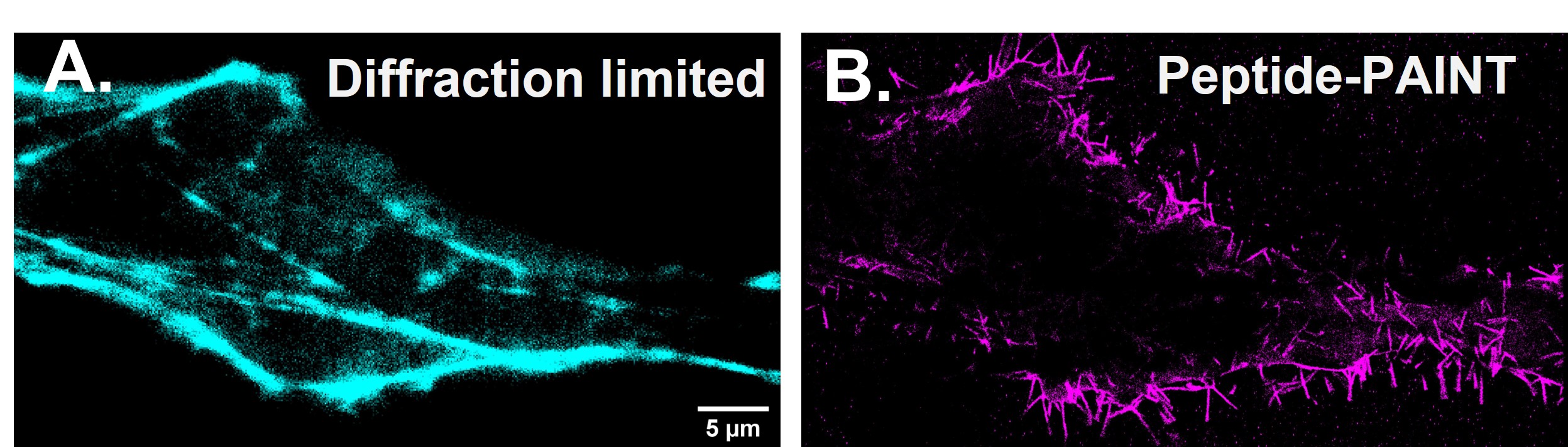

**Fig. S11: Peptide-PAINT imaging of neuroligin in live HeLa cells.** HeLa cells were transiently transfected with BFP-Lifeact (Lifeact, a peptide binds to actin) and E22-neuroligin plasmids.

| **Sample** | **Frame number** | **Exposure time (ms)** | **Localization precision (nm) from Picasso** | **Mean photon counts from Picasso** | **Uncertainty (nm) from ThunderSTORM** | **Mean photon counts from ThunderSTORM** | **Laser power (W/cm^2^)** |
| --- | --- | --- | --- | --- | --- | --- | --- |
| Vimentin-GFP-E22 | 10000 | 100 | 15 | 614 | 24 | 807 | 36 |
| Actin-GFP+E22-Lifeact | 10000 | 100 | 10 | 1034 | 12 | 986 | 36 |
| MAP4-E22+αTubulin-GFP | 20000 | 100 | 13 | 586 | 23 | 517 | 36 |
| Golgi7-GFP-E22 | 10000 | 100 | 14 | 723 | 20 | 968 | 36 |
| H2B-GFP-E22 | 12000 | 100 | 15 | 662 | 25 | 809 | 36 |
| E22-GluA2 + aHomer-YFP | 8000 | 100 | 11 | 1325 | 20 | 1569 | 36 |
| E22-Neuroligin+Homer-mGeos | 5000 | 100 | 9 | 1453 | 18 | 1496 | 74 |

**Sequences:**

**CK19**: CKIAALKE KIAALKE KIAAL

**E22**: EIAALEKEIAALEKEIAALEKK

**Vimentin-GFP-E22**

MAAMSTRSVSSSSYRRMFGGPGTASRPSSSRSYVTTSTRTYSLGSALRPSTSRSLYASSPGGVYATRSSAVRLRSSVPGVRLLQDSVDFSLADAINTEFKNTRTNEKVELQELNDRFANYIDKVRFLEQQNKILLAELEQLKGQGKSRLGDLYEEEMRELRRQVDQLTNDKARVEVERDNLAEDIMRLREKLQEEMLQREEAENTLQSFRQDVDNASLARLDLERKVESLQEEIAFLKKLHEEEIQELQAQIQEQHVQIDVDVSKPDLTAALRDVRQQYESVAAKNLQEAEEWYKSKFADLSEAANRNNDALRQAKQESTEYRRQVQSLTCEVDALKGTNESLERQMREMEENFAVEAANYQDTIGRLQDEIQNMKEEMARHLREYQDLLNVKMALDIEIATYRKLLEGEESRISLPLPNFSSLNLRETNLDSLPLVDTHSKRTLLIKTVETRDGQVINETSQHHDDLEGDPPVATMVSKGEELFTGVVPILVELDGDVNGHKFSVSGEGEGDATYGKLTLKFICTTGKLPVPWPTLVTTLTYGVQCFSRYPDHMKQHDFFKSAMPEGYVQERTIFFKDDGNYKTRAEVKFEGDTLVNRIELKGIDFKEDGNILGHKLEYNYNSHNVYIMADKQKNGIKVNFKIRHNIEDGSVQLADHYQQNTPIGDGPVLLPDNHYLSTQSALSKDPNEKRDHMVLLEFVTAAGITLGMDELYKAGA**EIAALEKEIAALEKEIAALEKK**

**E22-Vimentin-GFP**

M**EIAALEKEIAALEKEIAALEKK**GGGAAMSTRSVSSSSYRRMFGGPGTASRPSSSRSYVTTSTRTYSLGSALRPSTSRSLYASSPGGVYATRSSAVRLRSSVPGVRLLQDSVDFSLADAINTEFKNTRTNEKVELQELNDRFANYIDKVRFLEQQNKILLAELEQLKGQGKSRLGDLYEEEMRELRRQVDQLTNDKARVEVERDNLAEDIMRLREKLQEEMLQREEAENTLQSFRQDVDNASLARLDLERKVESLQEEIAFLKKLHEEEIQELQAQIQEQHVQIDVDVSKPDLTAALRDVRQQYESVAAKNLQEAEEWYKSKFADLSEAANRNNDALRQAKQESTEYRRQVQSLTCEVDALKGTNESLERQMREMEENFAVEAANYQDTIGRLQDEIQNMKEEMARHLREYQDLLNVKMALDIEIATYRKLLEGEESRISLPLPNFSSLNLRETNLDSLPLVDTHSKRTLLIKTVETRDGQVINETSQHHDDLEGDPPVATMVSKGEELFTGVVPILVELDGDVNGHKFSVSGEGEGDATYGKLTLKFICTTGKLPVPWPTLVTTLTYGVQCFSRYPDHMKQHDFFKSAMPEGYVQERTIFFKDDGNYKTRAEVKFEGDTLVNRIELKGIDFKEDGNILGHKLEYNYNSHNVYIMADKQKNGIKVNFKIRHNIEDGSVQLADHYQQNTPIGDGPVLLPDNHYLSTQSALSKDPNEKRDHMVLLEFVTAAGITLGMDELYK

**Golgi-GFP-E22**

MRLREPLLSGSAAMPGASLQRACRLLVAVCALHLGVTLVYYLAGRDLSRLPQLVGVSTPLQGGSNSAAAIGQSSGELRTGGAKDPPVATMDNTEDMVSKGEELFTGVVPILVELDGDVNGHKFSVSGEGEGDATYGKLTLKFICTTGKLPVPWPTLVTTLTYGVQCFSRYPDHMKQHDFFKSAMPEGYVQERTIFFKDDGNYKTRAEVKFEGDTLVNRIELKGIDFKEDGNILGHKLEYNYNSHNVYIMADKQKNGIKVNFKIRHNIEDGSVQLADHYQQNTPIGDGPVLLPDNHYLSTQSALSKDPNEKRDHMVLLEFVTAAGITLGMDELYKSAGA**EIAALEKEIAALEKEIAALEKK**

**H2B-GFP-E22** MPEPAKSAPAPKKGSKKAVTKAQKKGGKKRKRSRKESYSIYVYKVLKQVHPDTGISSKAMGIMNSFVNDIFERIAGEASRLAHYNKRSTITSREIQTAVRLLLPGELAKHAVSEGTKAITKYTSAKDPPVATMVSKGEELFTGVVPILVELDGDVNGHKFSVSGEGEGDATYGKLTLKFICTTGKLPVPWPTLVTTLTYGVQCFSRYPDHMKQHDFFKSAMPEGYVQERTIFFKDDGNYKTRAEVKFEGDTLVNRIELKGIDFKEDGNILGHKLEYNYNSHNVYIMADKQKNGIKVNFKIRHNIEDGSVQLADHYQQNTPIGDGPVLLPDNHYLSTQSALSKDPNEKRDHMVLLEFVTAAGITLGMDELYKSAGA**EIAALEKEIAALEKEIAALEKK**

**Lifeact-E22**

MGVADLIKKFESISKEEGDPPVATMA**EIAALEKEIAALEKEIAALEKK**

**MAP4-E22**

MVSRQEEAKAAVGVTGNDITTPPNKEPPPSPEKKAKPLATTQPAKTSTSKAKTQPTSLPKQPAPTTSGGLNKKPMSLASGSVPAAPHKRPAAATATARPSTLPARDVKPKPITEAKVAEKRTSPSKPSSAPALKPGPKTTPTVSKATSPSTLVSTGPSSRSPATTLPKRPTSIKTEGKPADVKRMTAKSASADLSRSKTTSASSVKRNTTPTGAAPPAGMTSTRVKPMSAPSRSSGALSVDKKPTSTKPSSSAPRVSRLATTVSAPDLKSVRSKVGSTENIKHQPGGGRAKVEKKTEAATTAGKPEPNAVTKAAGSIASAQKPPAGKVQIVSKKVSYSHIQSKCVSKDNIKHVPGCGNVQIQNKKVDISKVSSKCGSKANIKHKPGGGDVKIESQKLNFKEKAQAKVGGGFADPPVATMSVIKPDMA**EIAALEKEIAALEKEIAALEKK**

**αTubulin-GFP**

MVSKGEELFTGVVPILVELDGDVNGHKFSVSGEGEGDATYGKLTLKFICTTGKLPVPWPTLVTTLTYGVQCFSRYPDHMKQHDFFKSAMPEGYVQERTIFFKDDGNYKTRAEVKFEGDTLVNRIELKGIDFKEDGNILGHKLEYNYNSHNVYIMADKQKNGIKVNFKIRHNIEDGSVQLADHYQQNTPIGDGPVLLPDNHYLSTQSALSKDPNEKRDHMVLLEFVTAAGITLGMDELYKSRVRECISIHVGQAGVQIGNACWELYCLEHGIQPDGQMPSDKTIGGGDDSFNTFFSETGAGKHVPRAVFVDLEPTVIDEVRTGTYRQLFHPEQLITGKEDAANNYARGHYTIGKEIIDLVLDRIRKLADQCTGLQGFLVFHSFGGGTGSGFTSLLMERLSVDYGKKSKLEFSIYPAPQVSTAVVEPYNSILTTHTTLEHSDCAFMVDNEAIYDICRRNLDIERPTYTNLNRLISQIVSSITASLRFDGALNVDLTEFQTNLVPYPRIHFPLATYAPVISAEKAYHEQLSVAEITNACFEPANQMVKCDPRHGKYMACCLLYRGDVVPKDVNAAIATIKTKRSIQFVDWCPTGFKVGINYQPPTVVPGGDLAKVQRAVCMLSNTTAIAEAWARLDHKFDLMYAKRAFVHWYVGEGMEEGEFSEAREDMAALEKDYEEVGVDSVEGEGEEEGEEY

**E22-GluA2**

MQKIMHISVLLSPVLWGLIFGVSSARG**EIAALEKEIAALEKEIAALEKK**GGAGLAHMGGGGSGGGGSTGILLEASNSIQIGGLFPRGADQEYSAFRVGMVQFSTSEFRLTPHIDNLEVANSFAVTNAFCSQFSRGVYAIFGFYDKKSVNTITSFCGTLHVSFITPSFPTDGTHPFVIQMRPDLKGALLSLIEYYQWDKFAYLYDSDRGLSTLQAVLDSAAEKKWQVTAINVGNINNDKKDETYRSLFQDLELKKERRVILDCERDKVNDIVDQVITIGKHVKGYHYIIANLGFTDGDLLKIQFGGANVSGFQIVDYDDSLVSKFIERWSTLEEKEYPGAHTATIKYTSALTYDAVQVMTEAFRNLRKQRIEISRRGNAGDCLANPAVPWGQGVEIERALKQVQVEGLSGNIKFDQNGKRINYTINIMELKTNGPRKIGYWSEVDKMVVTLTELPSGNDTSGLENKTVVVTTILESPYVMMKKNHEMLEGNERYEGYCVDLAAEIAKHCGFKYKLTIVGDGKYGARDADTKIWNGMVGELVYGKADIAIAPLTITLVREEVIDFSKPFMSLGISIMIKKPQKSKPGVFSFLDPLAYEIWMCIVFAYIGVSVVLFLVSRFSPYEWHTEEFEDGRETQSSESTNEFGIFNSLWFSLGAFMRQGCDISPRSLSGRIVGGVWWFFTLIIISSYTANLAAFLTVERMVSPIESAEDLSKQTEIAYGTLDSGSTKEFFRRSKIAVFDKMWTYMRSAEPSVFVRTTAEGVARVRKSKGKYAYLLESTMNEYIEQRKPCDTMKVGGNLDSKGYGIATPKGSSLGTPVNLAVLKLSEQGVLDKLKNKWWYDKGECGAKDSGSKEKTSALSLSNVAGVFYILVGGLGLAMLVALIEFCYKSRAEAKRMKVAKNPQNINPSSSQNSQNFATYKEGYNVYGIESVKI

**E22-Neuroligin**

MALPRCMWPNYVWRAMMACVVHRGSGAPLTLCLLGCLLQTFHVLSQKGGGGS**EIAALEKEIAALEKEIAALEKK**GGGGSLDDVDPLVTTNFGKIRGIKKELNNEILGPVIQFLGVPYAAPPTGEHRFQPPEPPSPWSDIRNATQFAPVCPQNIIDGRLPEVMLPVWFTNNLDVVSSYVQDQSEDCLYLNIYVPTEDDIRDSGGPKPVMVYIHGGSYMEGTGNLYDGSVLASYGNVIVITVNYRLGVLGFLSTGDQAAKGNYGLLDLIQALRWTSENIGFFGGDPLRITVFGSGAGGSCVNLLTLSHYSEGLFQRAIAQSGTALSSWAVSFQPAKYARILATKVGCNVSDTVELVECLQKKPYKELVDQDVQPARYHIAFGPVIDGDVIPDDPQILMEQGEFLNYDIMLGVNQGEGLKFVENIVDSDDGVSASDFDFAVSNFVDNLYGYPEGKDVLRETIKFMYTDWADRHNPETRRKTLLALFTDHQWVAPAVATADLHSNFGSPTYFYAFYHHCQTDQVPAWADAAHGDEVPYVLGIPMIGPTELFPCNFSKNDVMLSAVVMTYWTNFAKTGDPNQPVPQDTKFIHTKPNRFEEVAWTRYSQKDQLYLHIGLKPRVKEHYRANKVNLWLELVPHLHNLNDISQYTSTTTKVPSTDITLRPTRKNSTPVTSAFPTAKQDDPKQQPSPFSVDQRDYSTELSVTIAVGASLLFLNILAFAALYYKKDKRRHDVHRRCSPQRTTTNDLTHAPEEEIMSLQMKHTDLDHECESIHPHEVVLRTACPPDYTLAMRRSPDDIPLMTPNTITMIPNTIPGIQPLHTFNTFTGGQNNTLPHPHPHPHSHSTTRV

**mGeos-Homer**

MSAIKPDMKIKLRMEGNVNGHHFVIDGDGTGKPFEGKQSMDLEVKEGGPLPFAFDILTTAFMYGNRVFAKYPDNIQDYFKQSFPKGYSWERSLTFEDGGICIARNDITMEGDTFYNKVRFYGTNFPANGPVMQKKTLKWEPSTEKMYVRDGVLTGDIHMALLLEGNAHYRCDFRTTYKAKEKGVKLPGYHFVDHCIEILSHDKDYNKVKLYEHAVAHSGLPDNARRSGGSGSGSSGGSSGSGGSGEQPIFSTRAHVFQIDPNTKKNWVPTSKHAVTVSYFYDSTRNVYRIISLDGSKAIINSTITPNMTFTKTSQKFGQWADSRANTVYGLGFSSEHHLSKFAEKFQEFKEAARLAKEKSQEKMELTSTPSQESAGGDLQSPLTPESINGTDDERTPDVTQNSEPRAEPAQNALPFSHSAGDRTQGLSHASSAISKHWEAELATLKGNNAKLTAALLESTANVKQWKQQLAAYQEEAERLHKRVTELECVSSQANAVHSHKTELSQTVQELEETLKVKEEEIERLKQEIDNARELQEQRDSLTQKLQEVEIRNKDLEGQLSELEQRLEKSQSEQDAFRSNLKTLLEILDGKIFELTELRDNLAKLLECS

**REFERENCES**

1.	Inavalli, V. V. G. K. *et al.* A super-resolution platform for correlative live single-molecule imaging and STED microscopy. *Nat. Methods* 1263–1268 (2019) doi:10.1038/s41592-019-0611-8.

2.	Schnitzbauer, J., Strauss, M. T., Schlichthaerle, T., Schueder, F. & Jungmann, R. Super-resolution microscopy with DNA-PAINT. *Nat. Protoc.* **12**, 1198–1228 (2017).

3.	Ovesný, M., Křížek, P., Borkovec, J., Švindrych, Z. & Hagen, G. M. ThunderSTORM: A comprehensive ImageJ plug-in for PALM and STORM data analysis and super-resolution imaging. *Bioinformatics* **30**, 2389–2390 (2014).

4.	Schindelin, J. *et al.* Fiji: An open-source platform for biological-image analysis. *Nat. Methods* **9**, 676–682 (2012).
